# Dynamic conformational states of apo and cabozantinib bound TAM kinases to differentiate active-inactive kinetic models

**DOI:** 10.1101/2021.04.21.440860

**Authors:** Gatta K R S Naresh, Lalitha Guruprasad

## Abstract

Tyro3, Axl, Mer (TAM) receptor tyrosine kinases (RTKs) are overexpressed in several human cancers. Cabozantinib, a small molecule inhibitor constrains the activity of TAM kinases at nanomolar concentrations. The dynamic active and inactive conformations of kinases play a crucial role in inhibitor binding and the activation of intracellular downstream signalling pathways. The all atom molecular dynamics (MD) simulations at microsecond timescale and longer provide robust insights into the structural details of conformational alterations of proteins due to their role cellular metabolic activities and signaling pathways. In this current study we report microsecond molecular dynamics (MD) simulations of apo, cabozantinib complexed active and inactive TAM RTKs and analysed the post-MD trajectories using the principal component analysis (PCA). Markov State Models (MSM) and transition pathways from Perron-cluster cluster analysis. For consensus, the 1µs atomistic simulations with enhanced computational algorithms indicated us to treat tyrosine kinase family by overwhelming dynamic states existence when bound to kinase inhibitors. The dynamic mechanistic pathways intrinsic to the kinase activity and protein conformational landscape in the TAM kinases are revealed due to the alterations in the P-loop, αC-helix, activation loop and αF-helix that result in breaking the regulatory and catalytic spines. We deciphered the long lived kinetic transition states of distinct active and inactive structural models from MD simulations trajectories of TAM RTKs bound inhibitor complex that have not been revealed so far.

## Introduction

Protein kinases are ubiquitous in all taxonomical classifications, and ∼ 2% genes in the human genome are associated with these cellular kinases.^1^ The human genome has ∼600 protein kinases, of which 90 proteins are encoded by tyrosine kinases and classified into receptor tyrosine kinases (RTKs) and non-receptor tyrosine kinases (NRTKs).^2a^ RTKs are single-pass membrane spanning protein receptors, recognised by extracellular ligands such as growth factors and hormones, which is a prerequisite for fundamental cellular processes. All RTKs share similar protein architecture; the N-terminal glycosylated extracellular region is stabilised by disulfide bonds followed by a membrane spanning α-helix, and an intracellular region comprising the kinase domain. The cell surface receptors are specific to extracellular ligands, the binding of a ligand causes receptor dimerization followed by kinase activation and intracellular autophosphorylation. This leads to a cascade of downstream signalling events such that the signalling pathways communicate within intracellular components and integrate with the nucleus. The activation of RTKs play vital role in the control of protein expression to regulate the normal physiological events in cell survival, proliferation, growth and death. The upregulation and overexpression of the mutated RTKs result in protein conformational changes and activity, leading to abnormal signalling pathways which have numerous effects on the role of proteins thus disrupting the regular rhythmic cellular functions. This often results in malignancy of cells that further develops and progresses into different human cancers^3^. The kinase domain in RTKs are universal drug targets for various diseases ranging from immunogenic, autoimmune, diabetes, heart disease and cellular cancers. These proteins are therefore important targets for cancer therapy, a leading cause of death worldwide.

The Tyro3, Axl and Mer (TAM) RTKs are a family of kinases, the extracellular N-terminal region comprises two immunoglobulin (Ig) domains, two fibronectin (FNIII) domains and the intracellular C-terminal region comprises the kinase domain. Extracellular ligands such as Growth arresting-specific 6 protein (Gas6) and protein S (Pros1) activate TAM kinases.^2b^ Gas6 has specific binding affinities for each of the three TAM RTKs, whereas Pros1 has specific affinity to bind Tyro3 and Mer.^4^ The three TAM kinase members have a high degree of sequence and structural homology in their kinase domains. The regular metabolic activity of TAM kinases requires extensive post-translational modifications such as glycosylation, ubiquitination and phosphorylation.^5^ Therefore, TAM RTKs consist of variable protein size 97, 98 and 110 kDa for Tyro3, Axl and Mer respectively, with molecular weights of 140-100 kDa for Mer, 165-140 kDa Axl and Tyro3 RTKs. The TAM RTKs play emerging roles in a variety of normal biological functions such as endothelial and vascular smooth-muscle homeostasis, spermatogenesis, bone physiology, controlling platelet aggregation.^6a^ TAM kinases are a class of innate immune checkpoints that participate in key steps of anti-tumoral immunity.^7^ TAM RTKs play crucial roles in disease conditions such as acute myeloid leukaemia, breast, colorectal, lung, ovarian cancers and glioblastoma. TAM RTKs are regulated by downstream signalling events that include cytosolic kinases such as JAK, p38, MEK and PI3K, these signalling pathways play essential role in cell growth, survival and apoptosis.^8^ The extracellular ligand, Gas6 promotes cell survival signalling pathway via active Axl to retain cardiac fibroblast cell types.^9, 10^

Kinase domains are viable drug targets for cancer treatment and several inhibitors have been designed and validated as cancer drugs.^11^ Cabozantinib is a small molecule inhibitor that is targeted towards multiple kinases such as Axl, c-Met, VEGFR2, RET, KIT and FLT3, and it is an FDA-approved drug for advanced renal cell carcinoma, hepatocellular carcinoma and medullary thyroid cancer. This molecule is under phase III clinical trials for effective VEGFR-targeted therapy. Cabozantinib is reported to bind TAM kinases with high affinity at nanomolar concentrations.^12, 13^**^, and^** ^14^

The activation of a kinase is triggered by binding to ATP and Mg^2+^ that results in autophosphorylation to initiate the kinase activity for the transfer of γ phosphate group to tyrosine or serine/threonine containing protein target. The ATP binding cleft is located between the N- and C-terminal lobes, and at the hinge region site connecting the two lobes.^15^ Additional structurally important regions required for the activity of a kinase include, the P-loop located between the β1 and β2 strands within N-terminal lobe (amino acids residues 539-553, numbering as per Axl throughout the manuscript, unless otherwise mentioned) that has high Gly rich sequence region (GKTLGEGEFGAVMEG). The catalytic helix, αC-helix (576-591) in the N-terminal lobe comprises the essential amino acid (Glu585) with its side chain fluctuating between the active and inactive states of kinase. The distinction between active/inactive states is also based upon the αC- helical movement towards or away from the ATP binding site. The disordered activation loop (689-724) in the C-terminal lobe has altered conformational states that are variable among the kinase structures reported so far. The orientation of side chains in the DFG sequence motif (690-692) indicate whether the kinase domain is in active or inactive state. An ionic interaction between the side chains of D581(αC-helix) - and K695 (activation loop) is important in the kinases structure and allostery. The synchronous fluctuations in the P-loop, αC-helix and activation loop leads to spatial alteration in the shape of the enzyme active site pocket and internal structural features such as the inward/outward rotation of αC-helix and expansion of the activation loop. In addition to these distinct structural features, the RTKs have two kinds of active sites such as regulatory substrate site and catalytic active site that become available during allosteric competitive inhibitor binding pathways in the cellular signal transduction process. Structure analyses revealed the presence of two non-contiguous structural motifs termed regulatory (R) and catalytic (C) spines ^16–18^ that are required for stabilizing the protein in active state.

It would be interesting to study how these RTKs are responsible for causing numerous cancers and also to design their inhibitors to arrest the disease. One approach would be to obtain key insights into the spatial dynamics of TAM kinases from classical long range MD simulations. This will help us to understand the cellular mechanistic pathways of inhibitor binding to kinases that will prevent internal signaling by up-regulation or overexpression of RTKs. In this manuscript, we report the highly unstable conformational states in TAM RTK domains by studying the apo and cabozantinib bound active and inactive states of TAM RTKs each for 1 µs MD simulations using AMBER 18.14 suite of programs.

## Results

The three-dimensional structures of active and inactive forms of Axl kinase domain were taken from the crystal structure (5U6B, B and A chains) ^6b^ on the basis of which, the homology models of active and inactive forms of Mer and Tyro3 kinase domains were constructed and validated.^35c^ The amino acid sequence alignment of Tyro3, Axl, Mer kinases is shown in supplementary **Figure S1a**, and the three-dimensional structures of the active and inactive forms display significant conformational alterations in the P-loop, αC-helix and activation loop (**Figure 1)**. The C- spine and R- spine dictate the positions of ATP and substrate in the regular kinase protein active and inactive states, respectively. They play a key role in the catalysis of kinases while binding with ATP. We have mapped the location of R-spine and C-spine on the structures of TAM kinases based on the structures of C-Src.**^1^**^9a^ The R-spine consists of four different non-consecutive hydrophobic amino acid residues aligned vertically from N-terminal lobe towards the C-terminal lobe through the activation loop.^20a^ Selection of these hydrophobic residues in Axl kinase domain comprise β_4_-strand Leu620; αC-helix Met589; DFG motif Phe691; catalytic loop His670 and an additional Asp731 from the C-terminal lobe represent the R-spine across the kinase domain (**Figure 2 a-c**). The C-spine consists of eight different non-consecutive hydrophobic amino acid residues aligned vertically from N-terminal lobe towards the C-terminal lobe through the hinge region. Selection of these hydrophobic residues in Axl kinase domain comprise, V550 (β2-strand), A565 (β3-strand), F622, L628 (hinge region), M679, L680 (catalytic loop), M739, I742 (αF-helix from C-terminal lobe) (**Figure 3 a-c**) (**Figure S1b and supporting videos 0-Axl-apo, 1-Axl-Cabo-active and 2-Axl-Cabo-inactive**).

**Figure 1:**
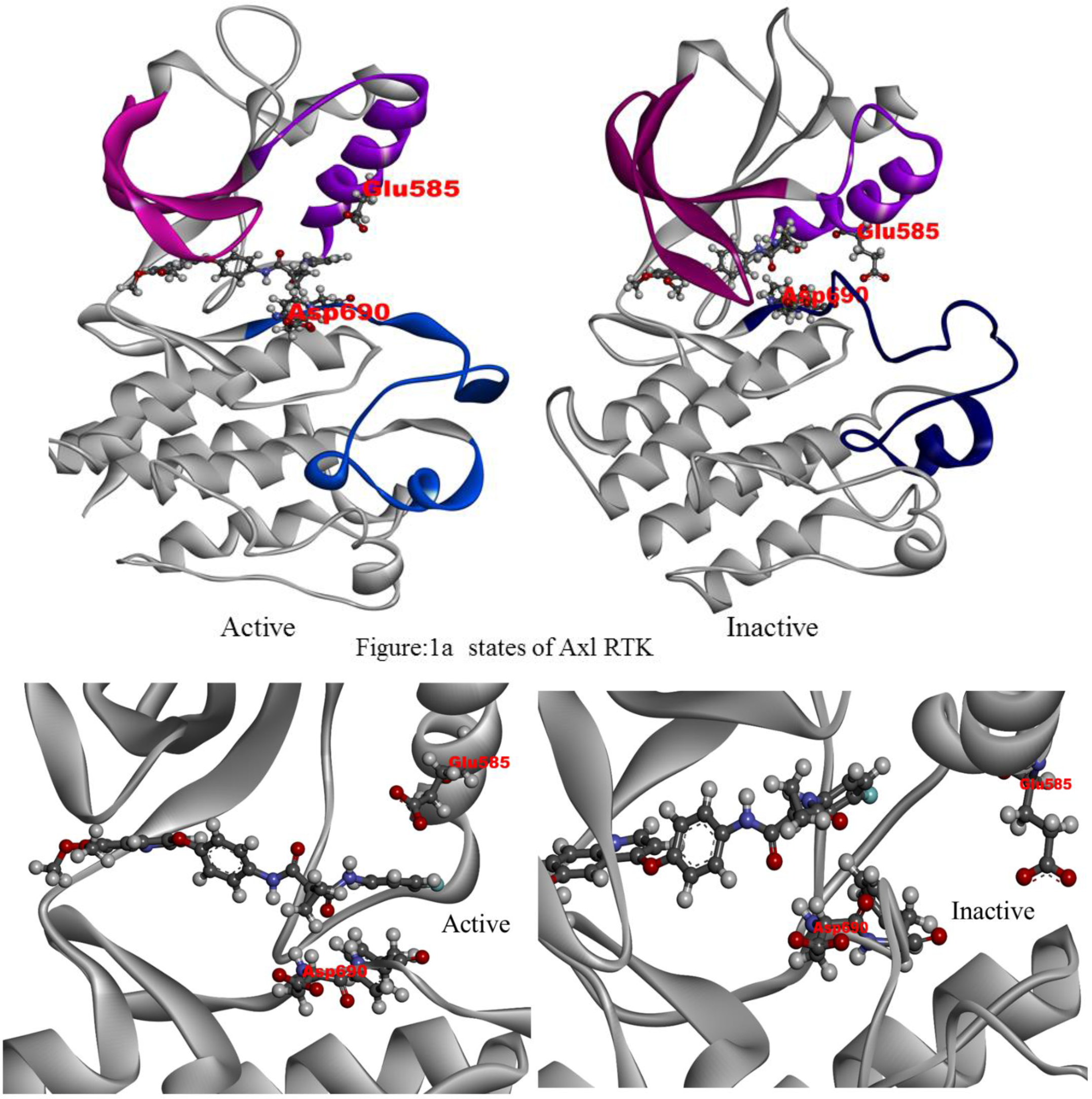
The active and inactive states of cabozantinib bound Axl receptor kinase. Axl (White); Cabozantinib (elemental color); Glu585-α C-helix - Asp690-Activation-loop.

**Figure 2:**
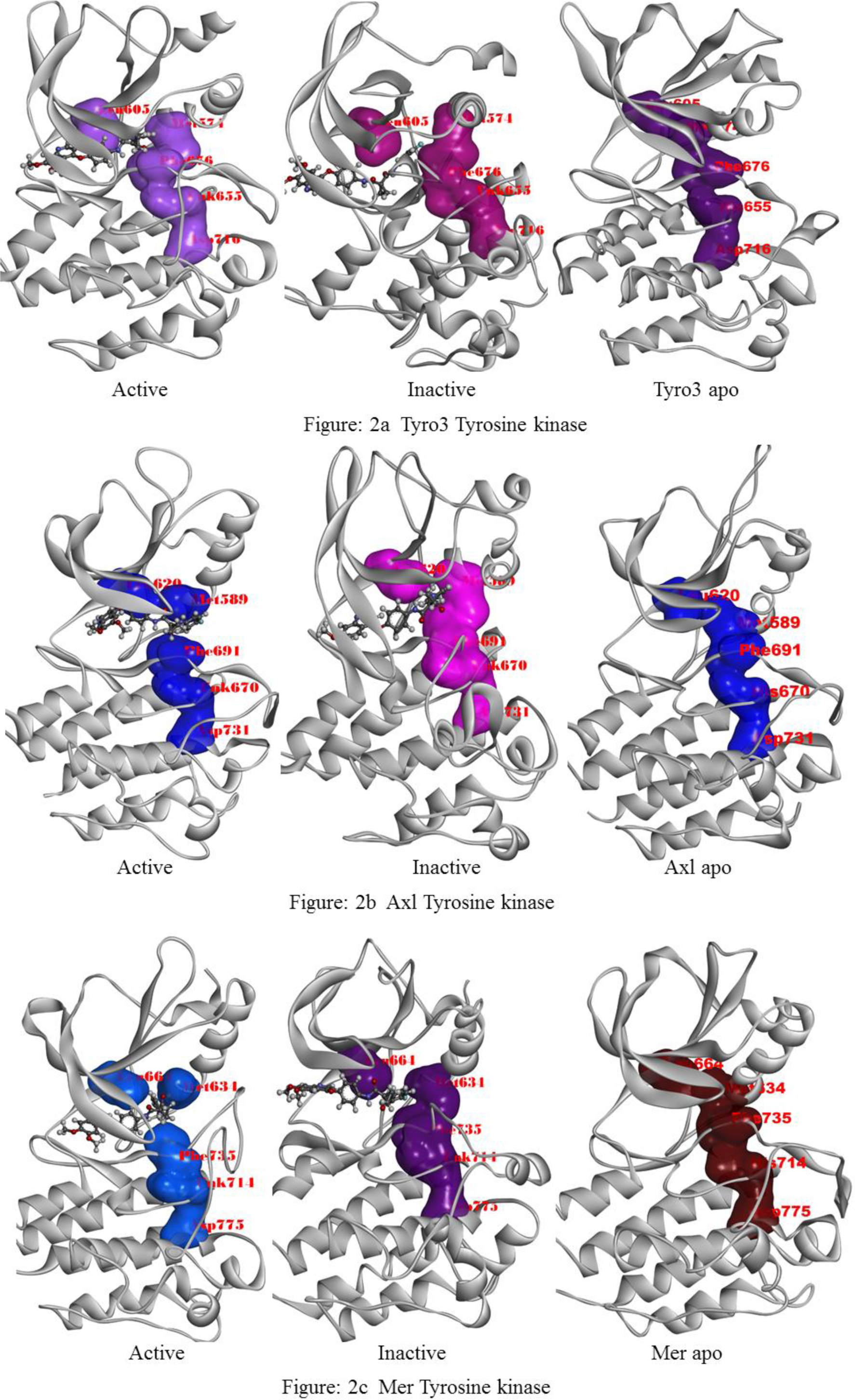
Regulatory spine (R-spine) analysis of TAM RTK’s – Cabozantinib bound active and inactive, apo models at 1μs. 2a) Tyro3; 2b) Axl; 2c) Mer;

**Figure 3:**
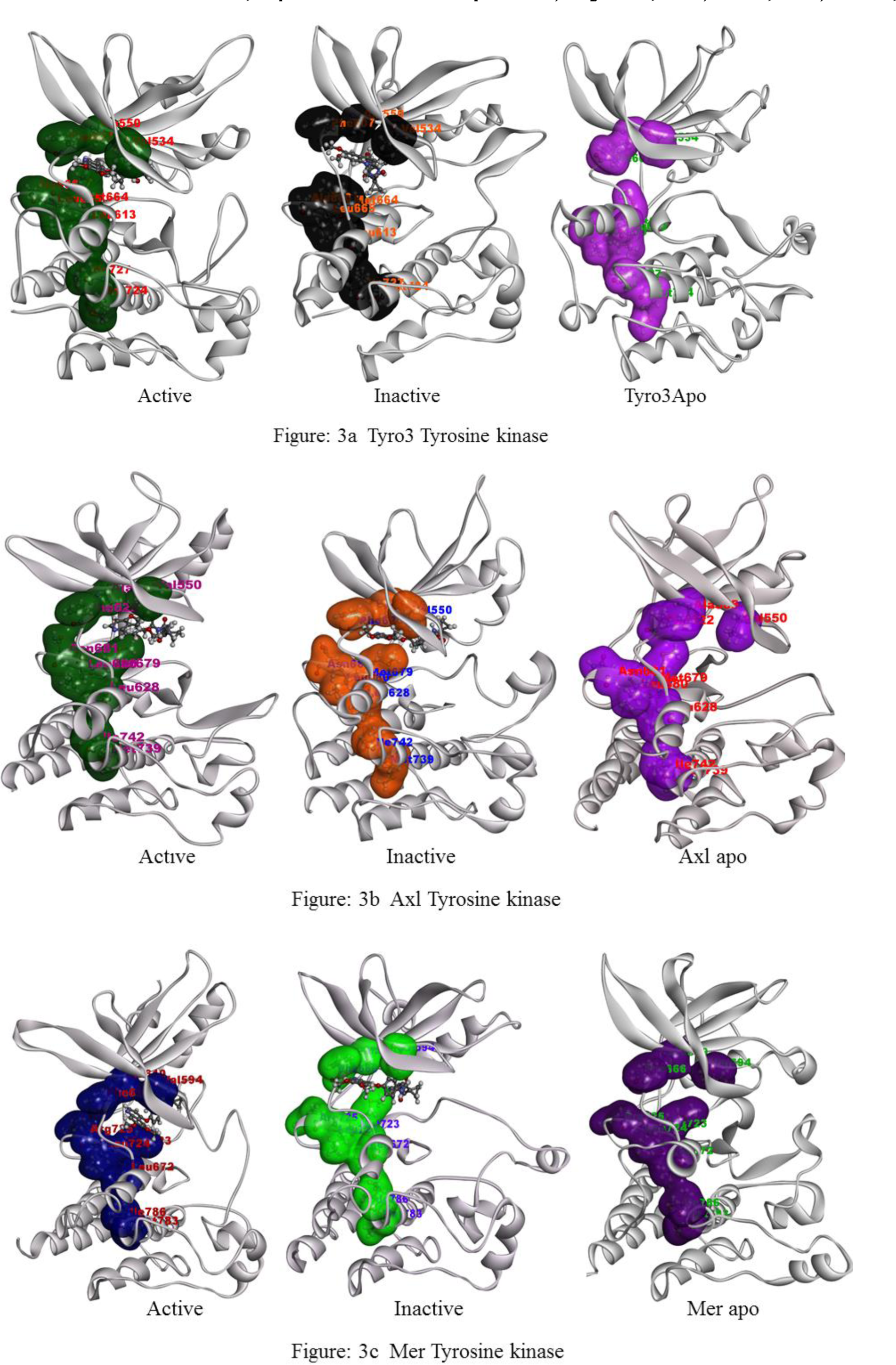
Catalytic spine (C-spine) analysis of TAM RTK’s – Cabozantinib bound active and inactive, apo models at 1μs. 3a) Tyro3; 3b) Axl; 3c) Mer.

From the docking of cabozantinib into TAM kinases, we observed that it binds to the ATP binding pocket mediated via several non-bonding interactions as described earlier.**^3^**^5d^ The hinge region residues F622, M623 (Axl) interact with nitrogen contained dimethoxy quinoline ring of cabozantinib. The para-fluoro phenyl interact with F691 aromatic ring (DFG motif in Axl) and D690 is also form hydrogen bond with amide linkage contained cyclopropyl ring of inhibitor. The structures of apo, active and inactive TAM kinases complexed with cabozantinib were subjected to 1 µs MD simulations each using AMBER. From the RMSD plots (**Figure 4a**), it can be seen that the structures converged at about 50 ns of MD simulations and the RMSD values lie within a narrow range from 2-4.5 Å. The apo Tyro3 and Axl have relatively higher RMSD values compared to the active and inactive cabozantinib complexes. Active Axl-cabozantinib complex is the most stable complex among all the systems studied. The Axl kinase bound cabozantinib shows lower variations in the active and inactive states when compared with the apo form. Similarly, apo Tyro3 has higher RMSD compared to the active and inactive states complexed with cabozantinib. However, all the three systems of Mer kinase have almost similar RMSD indicating that Mer RTKs have close associates of these states during longer time scales of MD simulations. The RMSD analysis of specified regions in kinases are the key components to describe the distribution among inactive and active states. The RMSD in the N-terminal P-loop is nearly similar in all the nine molecular systems, indicating that the structures consist of inseparable states due to close association in all molecular systems studied. The RMSD values of αC-helix region is distinguished among all TAM kinases. The active and apo states of Tyro3, the inactive and apo states of Axl, and the apo Mer kinase have higher and nearly similar RMSD values among all the kinase states. The RMSD is lowest in the inactive Tyro3, Axl active, active and inactive Mer complexes. The RMSD values of the αC-helix region is quite opposite to the activation loop (**Figure 4d-4f**). The active state Tyro3, active and apo states of Axl, and apo state of Mer kinase have lower and nearly similar RMSD values among all states. The activation loop in the apo and inactive Tyro3, inactive Axl, active and inactive Mer has highly dynamic conformation as can be seen from the higher RMSD values. Among all the states studied, the inactive Axl activation loop is highly variable. The above RMSD results are further confirmed by RMSF plots of all the TAM RTK molecular systems studied. The RMSF values of apo, active and inactive states of TAM kinases is shown in the **Figure 5**. From the analyses of the RMSD and RMSF plots, we believe that TAM kinase domains have unique hidden dynamic states that can be distinguished from further analyses of MD trajectories.

**Figure 4:**
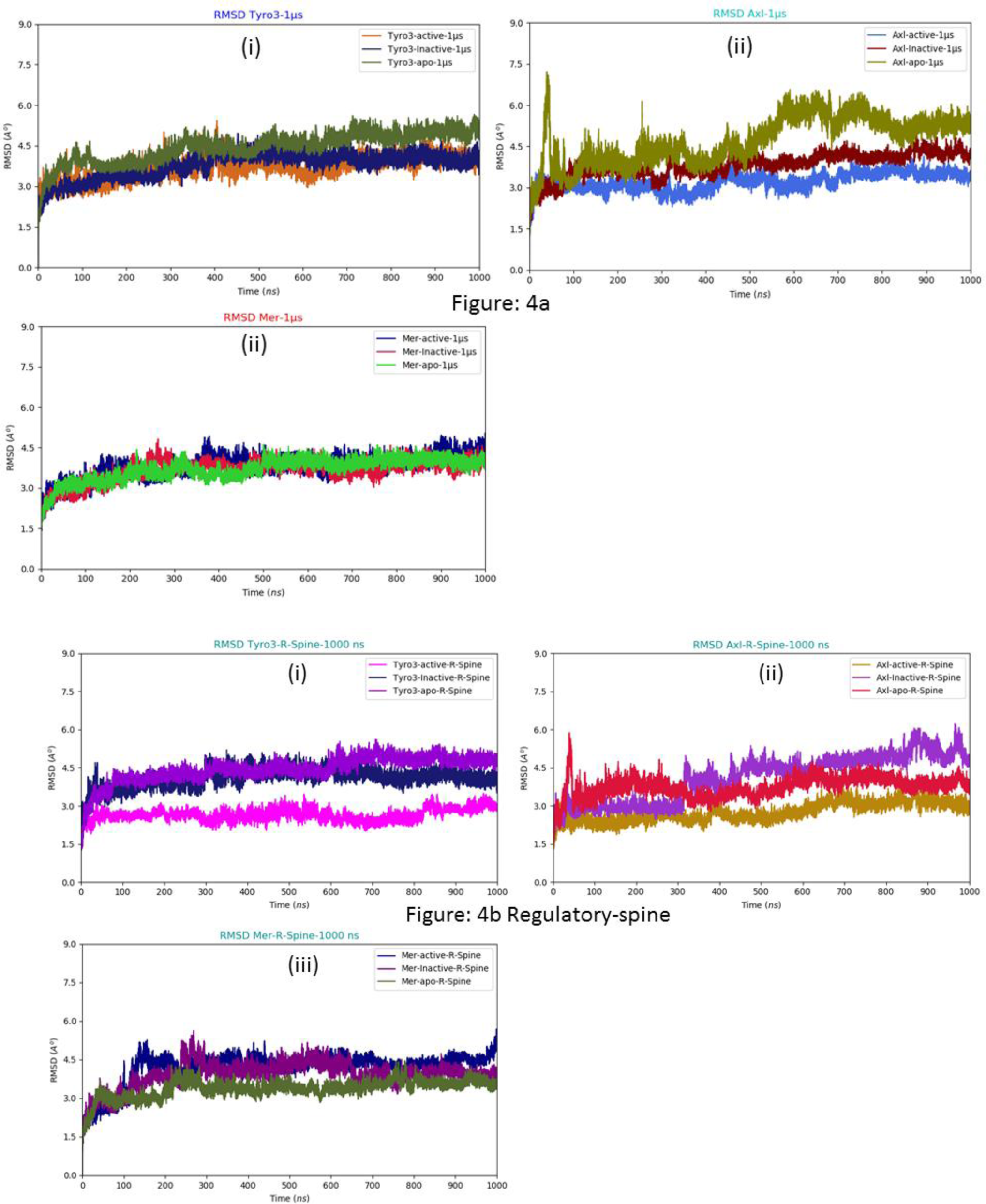
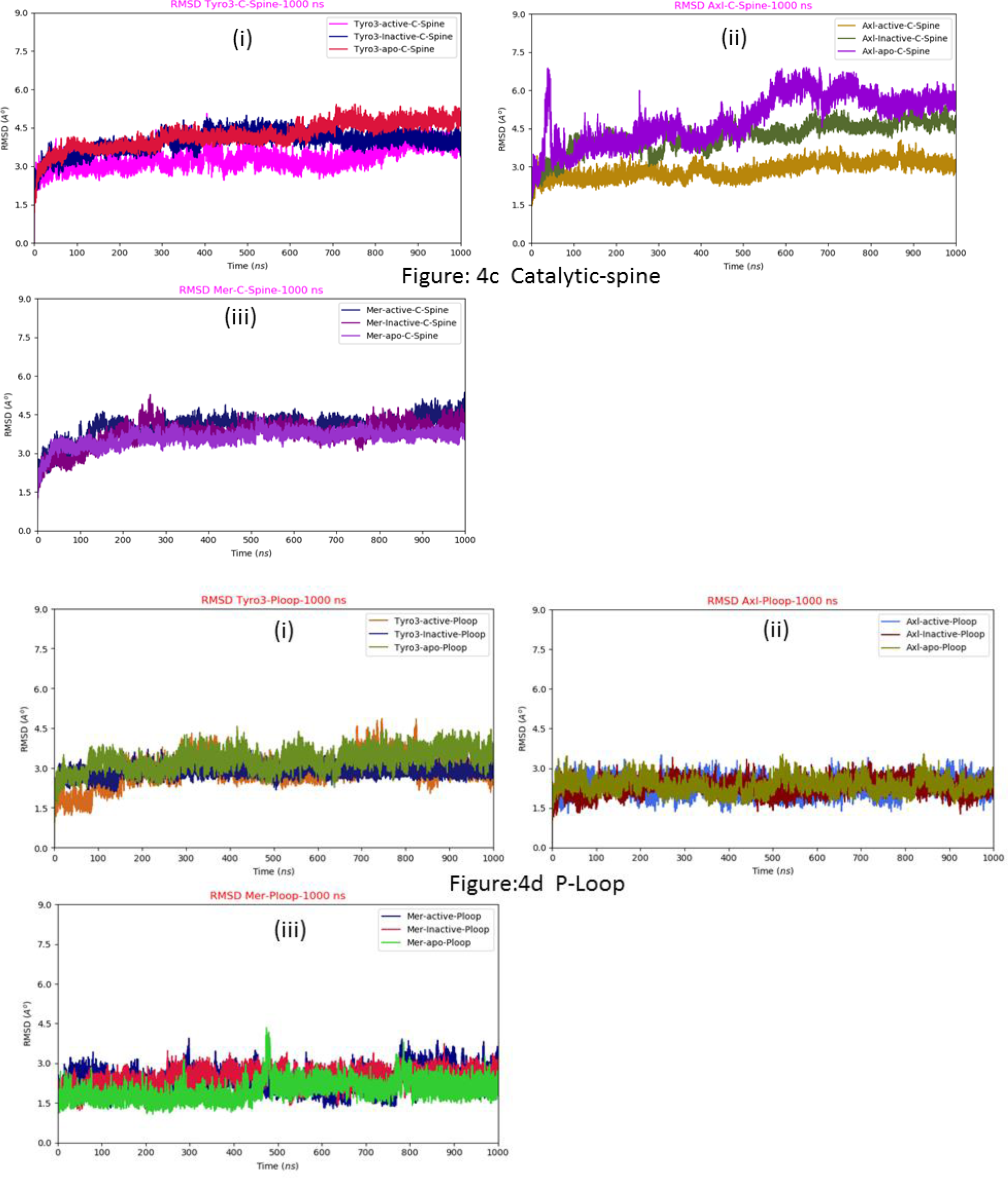
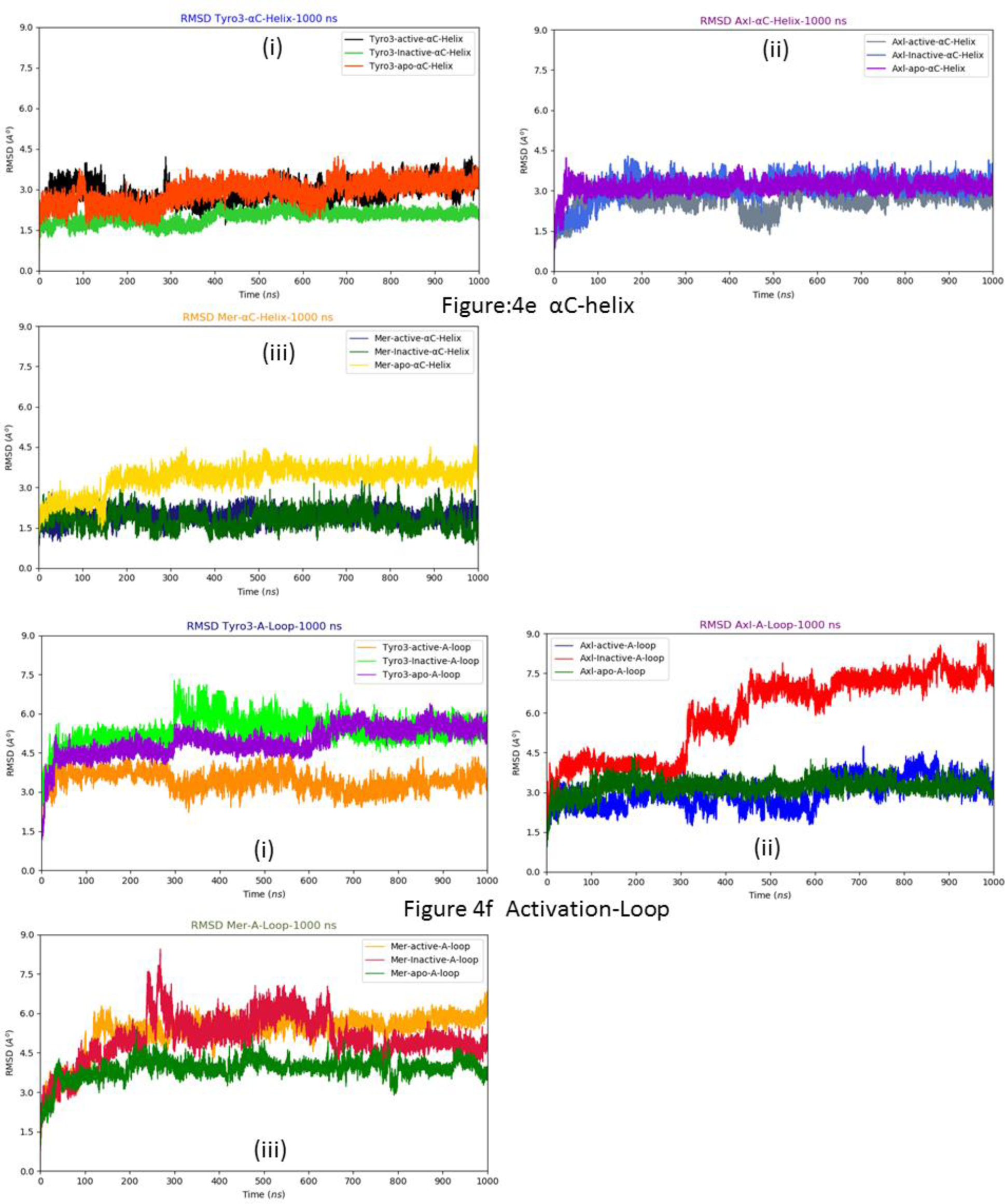
RMSD plots of apo and cabozantinib bound active and inactive TAM RTKs from 1μs MD simulations. (4a) Tyro3 Axl Mer proteins (4b) R-spine (4c) C-spine (4d) P-loop (4e) αC-helix (4f) Activation-loop.

**Figure 5:**
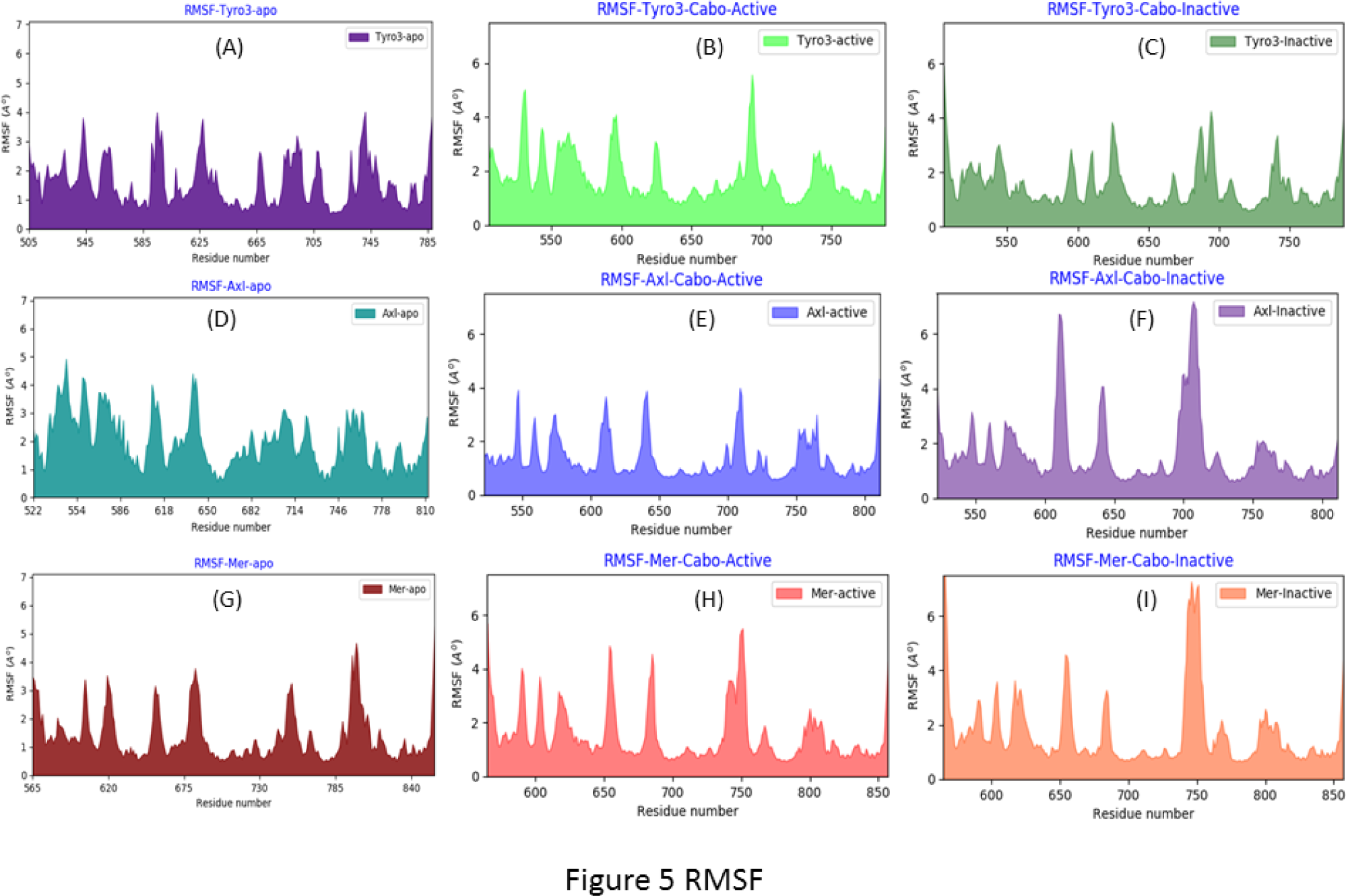
RMSF plots of TAM RTKs from 1μs MD simulations. In this figure, for the sake of convenience all RTK kinases are numbered from Tyro3-505; Axl-522; Mer-565 onwards. Axl indexing 539-553 (β1-β3 turn in the N-Terminal domain-P-loop); 579-591 (αC-Helix in the C-Terminal domain); 689-724 (Activation-Loop);

### Active – inactive kinetic state models of TAM RTKs

From the three-dimensional crystal structures, homology models and MD simulations results of TAM RTKs, the kinetic state models are defined according the internal structural dynamical features such as P- loop (539–553), αC-helix (576 –591) and activation loop (689–724). The active/inactive conformers of TAM kinases are distinguished by the side chain orientations; in the active state, the side chain of Glu585 on αC-helix is rotated inwards towards the substrate binding site and the side chain of Asp690 from the DFG motif also projects towards the active site. The outward orientation of Glu585 side chain away from the substrate and Asp690 side chain inwards into kinase active site is indicative of an inactive state of kinase.^6c^ These are key structural features implicated in the regulation of protein kinase activity and influence the effective binding of inhibitors and the specificity of inhibitor binding to RTKs. The binding of cabozantinib influenced various states of active/inactive models in Tyro3, Axl and Mer kinase domains. The active kinetic models are shown as αC-helix inward rotation and activation loop extended to further maximize inhibitor binding site. In the inactive state, the αC-helix undergoes outward rotation, followed by the activation loop inward folding so as to minimize the drug binding active site that can be seen from **Figure 1**.

### TAM kinase-cabozantinib complex activation pathway

The kinetic states appear due to stereo-spatial arrangement of certain residues in specific α- helices, β-sheets and loop regions in the kinase domain. These kinetic states provide key insights into the activation of protein kinase in the presence of inhibitor bound to the active site. The apo, active/inactive Axl-cabozantinib molecular systems consist of well-defined kinetic state models during the MD simulations. The precise way of Axl kinase active and inactive states represented in local spatial pattern can be accessed via R-spine and C-spine. The R-spine controls substrate molecule in the active site (αC- helix and activation loop). The C-spine regulates catalysis by allowing the ATP binding site at hinge region. The inactive kinase state should be converted into active state with help of substrate binding at activation loop thorough the influence of R-spine hydrophobic residues which connect the dynamical movement of catalytic loop in αF-helix. It is the coordination between R-spine and C-spine to evolve a dynamical conformation for the transfer of γ-phosphate group to target residue from ATP molecule binding hinge residue.^19b, 20b, 21, 22^

The R- spine is continuous and linear in the case of normal metabolic kinase activity. The hydrophobic surface in the R-spine is vertically aligned (Leu-Met-Phe-His) in the apo form of all TAM kinases as can be seen from the **Figures (2a-c)**. In the Axl active state, the R-spine is broken in the inhibitor bound form due to the expansion of activation loop that results in the extended space between αC-helix- Met589 and DFG motif -Phe691 in the ATP binding site. The inactive Axl bound to cabozantinib has a broken R-spine due to the expansion of space between αC-helix Met589 and β_4_-strand Leu620 as a result of the outward rotation of αC-helix. The fragmentation of R-spine occurred in similar way in the cabozantinib bound Tyro3 RTK in the active and inactive states, due to the increased distance between P-loop and αC-helix. In the case of the active state Mer RTK, R-spine fragmentation occurs between the P-loop, αC-helix and activation loop, whereas in the inactive Mer, R-spine fragmentation occurs between P-loop and αC-helix.

In the active kinase state, the R-spine is broken in all TAM RTKs but C-spine is retained only in Axl and Mer RTKs with no breakage in the hinge region. In the inactive kinase state of TAM RTKs, both the spines are broken in a similar region. Therefore, the cabozantinib binding active state kinase influences at specified locations in the R-spine residues rather than C-spine. This can lead the C-spine to initiate catalytic activity towards passive mechanism to alert the body immune system with help of chemokines. Whereas, in the inactive kinase state, the inhibitor binding to the regulatory active site, R-spine activates either the dynamical movement of catalytic loop or C-spine to initiate the catalysis process with help of cofactor ATP. As a consequence, both the spines are completely broken to trigger apoptosis in malignant cells.

The analysis of salt bridge distance between the activation loop and αC-helix reveals the hidden conformers among apo, active and inactive states. The salt bridge interaction in apo kinase is retained between the residues Asp (Axl-581) or Glu (Mer-626, Tyro3-566) of αC-helix and Lys (Axl-695, Mer- 739) or Arg (Tyro3-680) of activation loop (A-loop), within a distance of (Tyro3-3.44 Å; Axl-3.99Å; Mer-3.86Å). The salt bridge distance between Asp/Glu (αC-helix) - Lys/Arg (A-loop) of active states increases in Axl and Mer RTKs due to the expanded core in the inhibitor binding site in RTKs (Tyro3-3.16 Å; Axl-13.92Å; Mer-10.28Å), but the inactive states of TAM RTKs salt bridge distance between αC-helix and activation loop is lower for Axl RTK (Tyro3-6.82 Å; Axl-2.83Å; Mer-8.83Å) (**Figure 6 a-d**). These salt bridge distances are indirect support to stationary state distribution in apo TAM RTKs. But the apo Tyro3 has unique kinetic transition occurring due its alterations in the state distribution. The salt bridge is broken in active states of TAM kinase more precisely than inactive state forms. This could happen due the state specificity. The salt bridge should be retained in the apo form but are stable in inactive states of Tyro3 and Axl rather than slightly broken in Mer inactive states. The salt bridge distance analysis provides clear evidence that the kinases co-exist in active and inactive state models while binding with inhibitor at the active site. The large distance across in the regulatory site of kinase active states occurred due to β-sheet formation in activation loop and inward rotation of αC-helix. This causes the extended nature of regulatory active site between αC-helix and activation loop. The inactive state models have αC-helix outward rotation and activation loop undergoes shift to helical loop to minimize the active space across αC-helix and C-lobe in the RTKs. These results are further supported from R-spine analysis. All TAM RTK apo forms have retained R-spine as a result of the retention of salt bridge interaction with unaltered distance between Asp/Glu (αC-helix) - Lys/Arg (A-loop). But most of the active states in Axl and Mer forms have broken R-spine between αC-helix and activation loop therefore the distance between these domains is extended and the salt bridge interaction is disturbed due to the increased distances between Asp/Glu (αC-helix) - Lys (activation-loop). In the inactive state of TAM RTKs, the R-spine is broken between P-loop and αC-helix, however, the salt bridge distances between Asp/Glu (αC-helix) - Lys/Arg (activation loop) residues are preserved.

**Figure 6:**
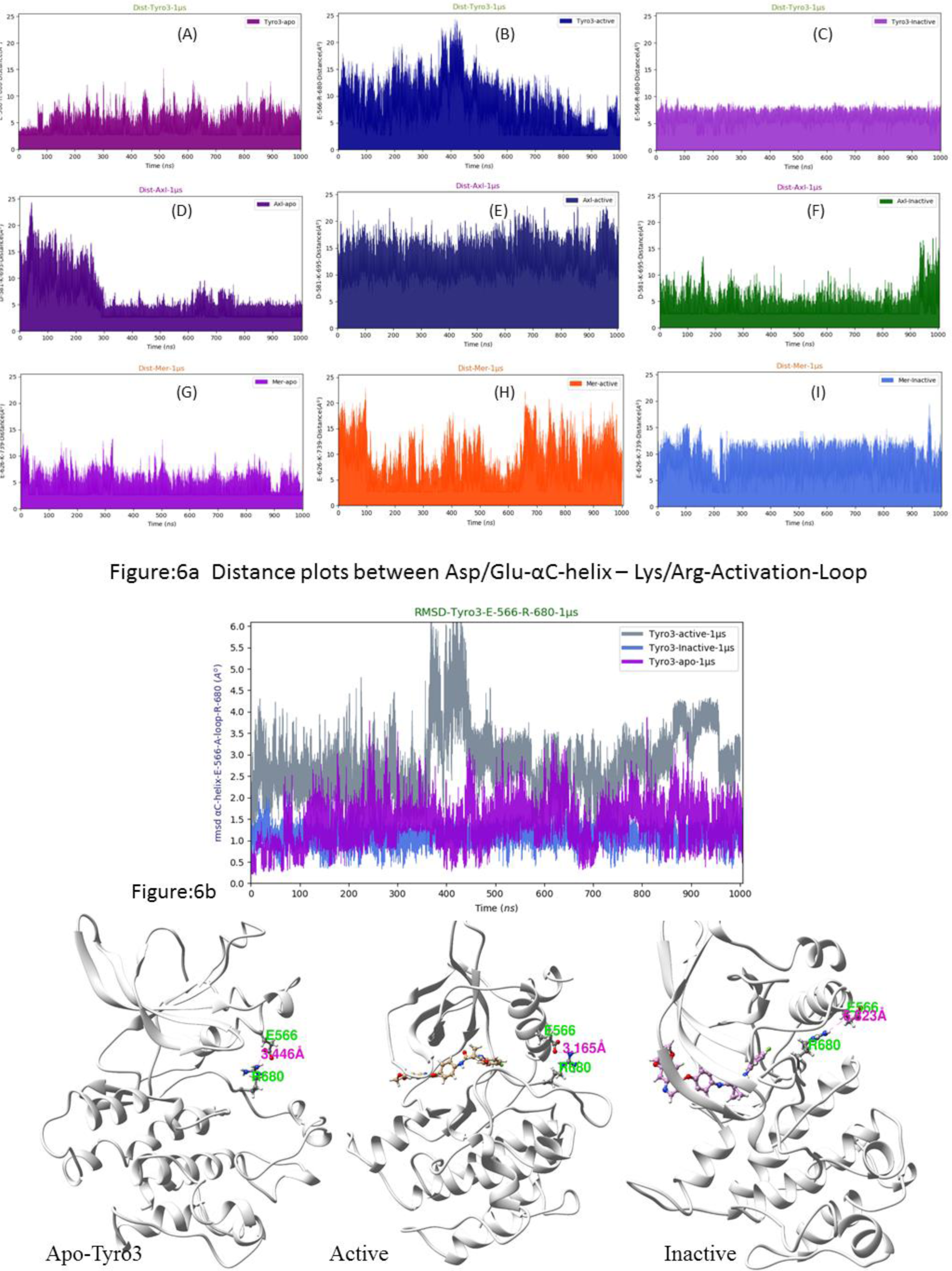
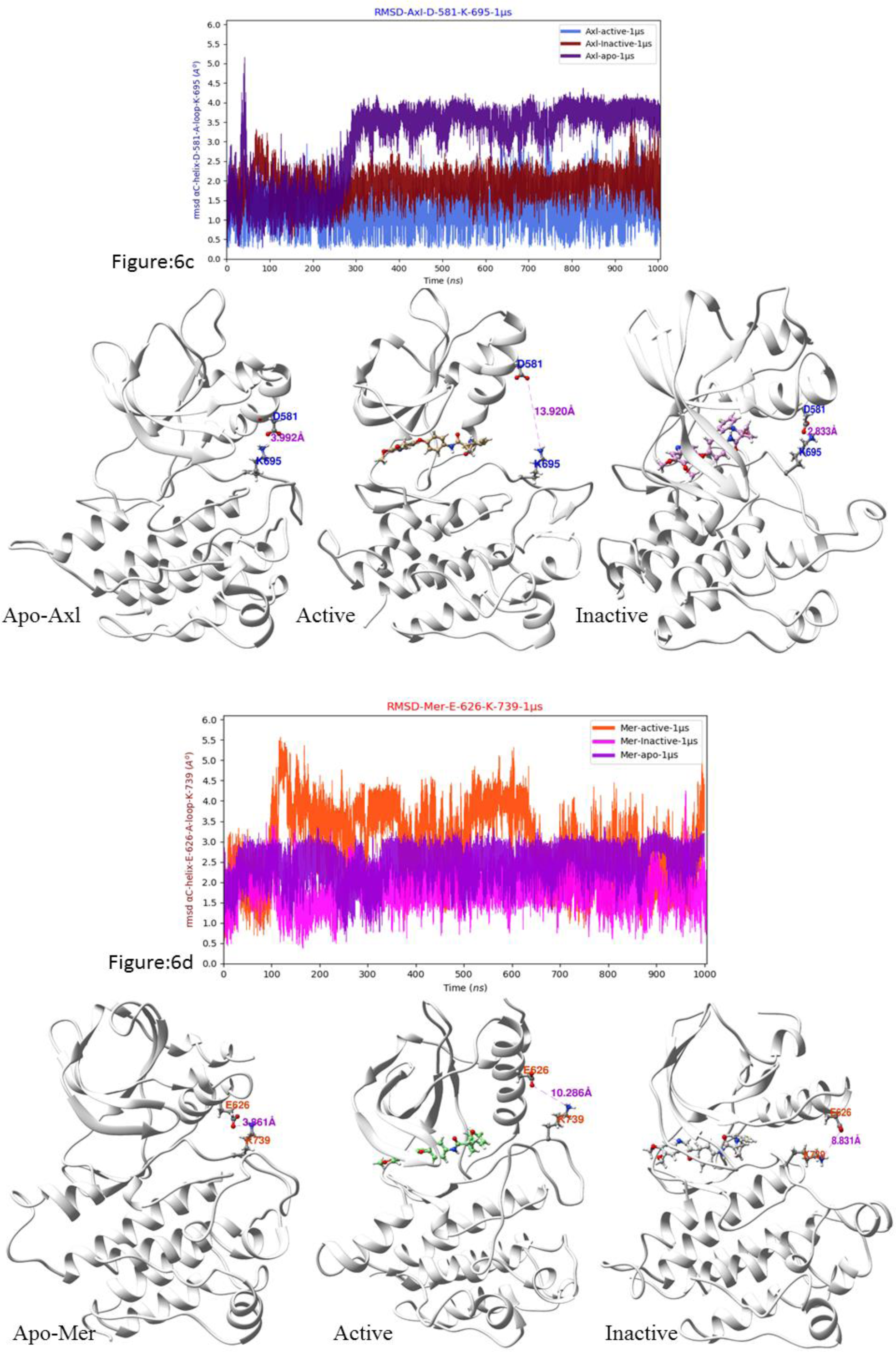
Distance plots between side chains of Asp/Glu-αC-helix – Lys/Arg- Activation-Loop pairs in apo and TAM-Cabozantinib bound active and inactive RTK from 1μs MD simulations. (6a) Column 1 - Apo; Column 2 - Active; column 3 – Inactive; (6b) Tyro3 (E566 – R680); (6c) Axl (D581 – K695); (6d) Mer (E626 - K739);

### Post-MD data analysis of TAM RTKs

The preliminary MD simulations data acquired from AMBER trajectories were analyzed to ensure kinetically active and inactive states were investigated with help of PCA. PCA analysis was carried out on 1K conformer samples of trajectories out of 40K for clear visualization of data points from kinetic transition states in the active to inactive kinases (**Figure 7 a-c**). The histogram showed that the random distribution of all nine kinase state trajectories data was extrapolated as training and test sets of individual components validated with shuffle-split cross-validation in PCA plot. All apo and inhibitor bound forms of TAM RTKs have random distribution of states that are very unique in nature from the respective scatter plots of kinase trajectory analysis. This is a preliminary analysis to propose the hidden dynamic states existing in longer MD timescale simulations and trajectory data of kinases.

**Figure 7:**
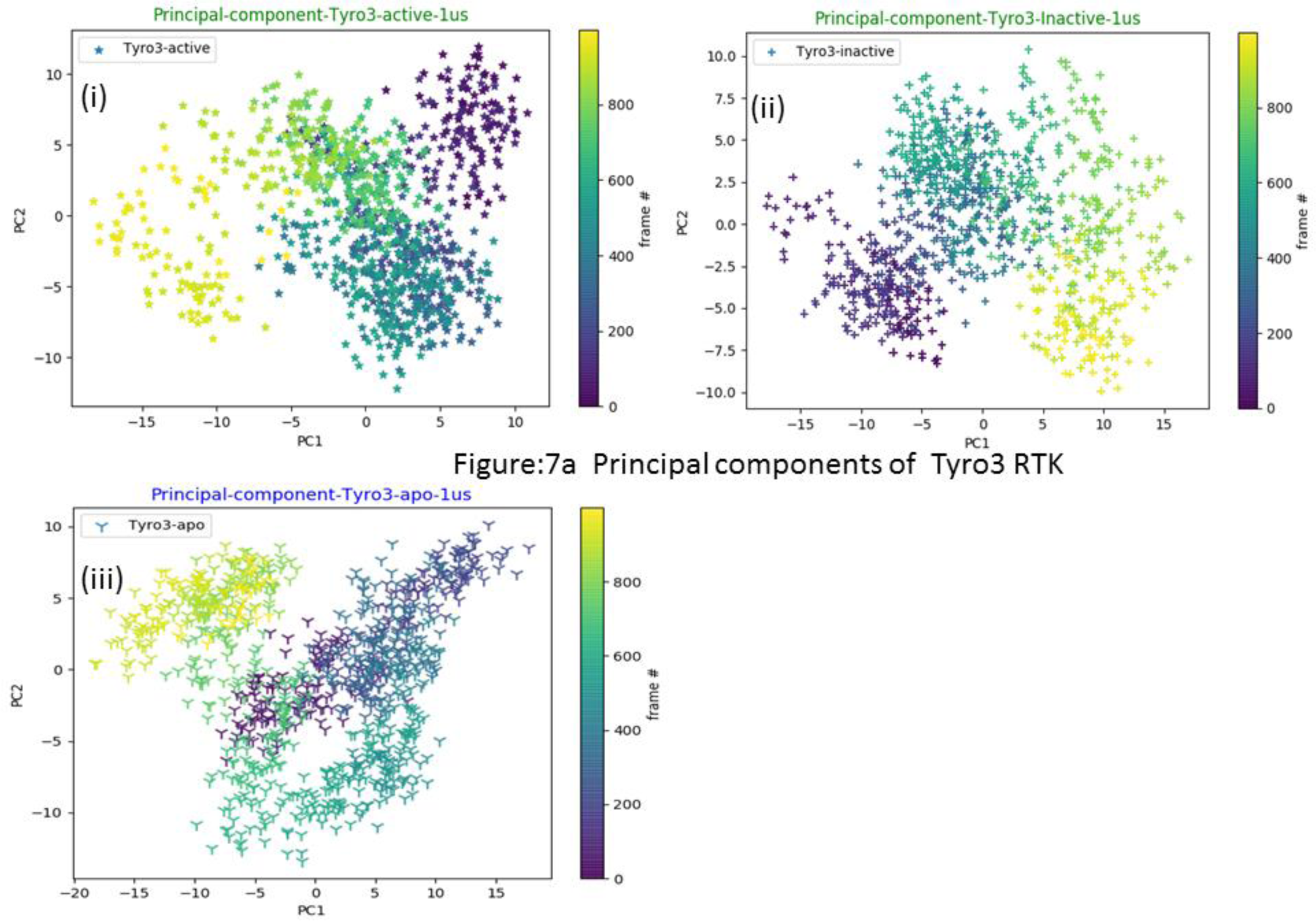
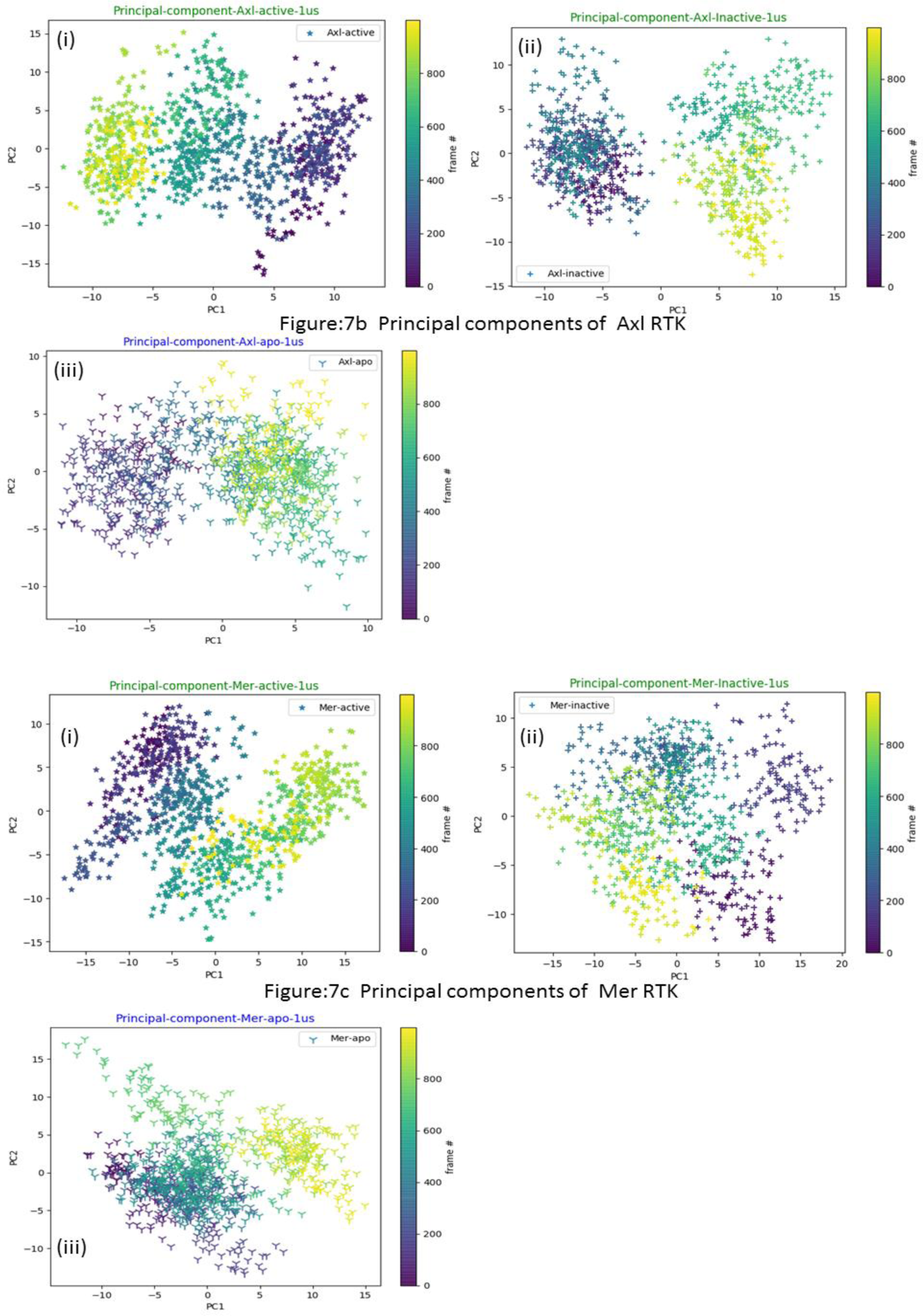
Principal component analysis of TAM kinase – apo & Cabozantinib bound active and inactive RTK from 1μs MD simulations. (i) Active; (ii) Inactive; (iii) Apo; 7a) Tyro3 (7b) Axl (7c) Mer

The metastable kinetic models were built based upon advanced trajectory data analysis using python based scripts. All TAM trajectories data was sampled into vectorized and clustering was done using keras-state algorithm for MSM model generation.^23,^ **^5^**^2b^ The MSM data of TAM kinases was bootstrapped from 1000 ns of trajectory data to generate Hidden Markov state models (HMM) to reveal the unfolding and refolding of the activation loop from active state to inactive states. The metastable trajectories are well converged as shown by VAMP score. Discrete clustering of protein backbone state distribution featurisation was performed to show distinct kinetic stable states in all TAM kinases. All HMM states are key intermediate conformers to describe the kinase inhibitory activity when bound to cabozantinib. As per the analysis of metastable kinetic state forms, higher numbers of active state models are present in Axl and Mer than the number of kinetic transitions states of inactive forms. However, the Tyro3 has approximately similar numbers of state models in their respective active and inactive states which are included in state distribution plots. The mean free passage time (MFPT) error bars were validated with Bayesian HMM model validation with lag time of 50 states (**Figure 8A-C**). From these analyses we infer that Tyro3 RTK states have combined and co-existed metastable state transitions among the active and inactive forms rather than the dominance of either active or inactive kinetic states as observed in Mer and Axl RTKs. Therefore, the Tyro3 has more intermediate states than Axl and Mer. The MFPT values of Tyro3 indicate that activation and deactivation happens in equal ratio (below 100 ns **Figure S2A-C**) whereas the Axl and Mer have different activation timescales (after 200 ns) and their deactivation takes place around 100 ns timescales (**Figure S2D-I**). The influence of these major changes in the kinase domain is due to the conversion of active to inactive states through kinetic transition metastable equilibrium states.

**Figure 8:**
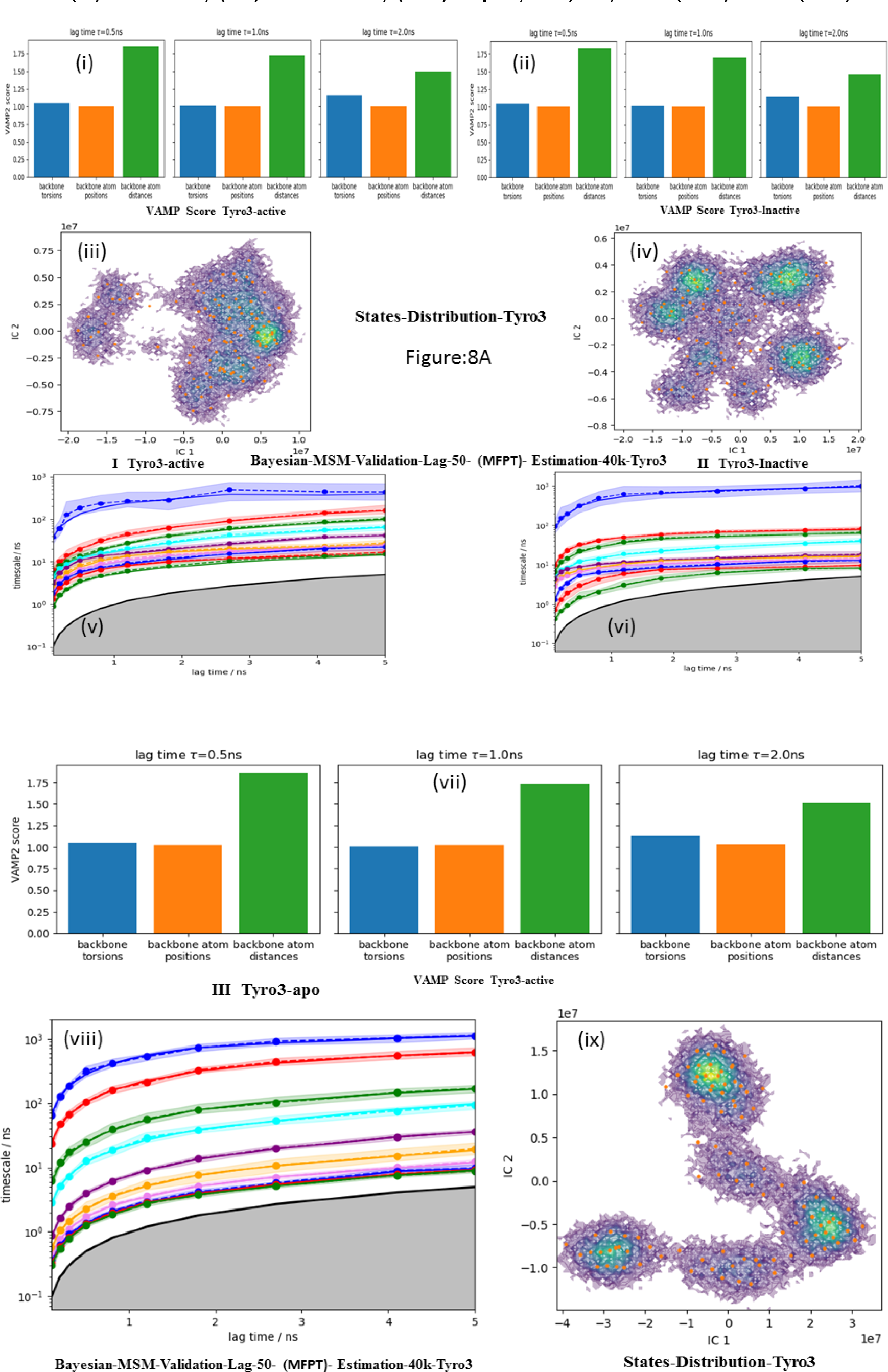
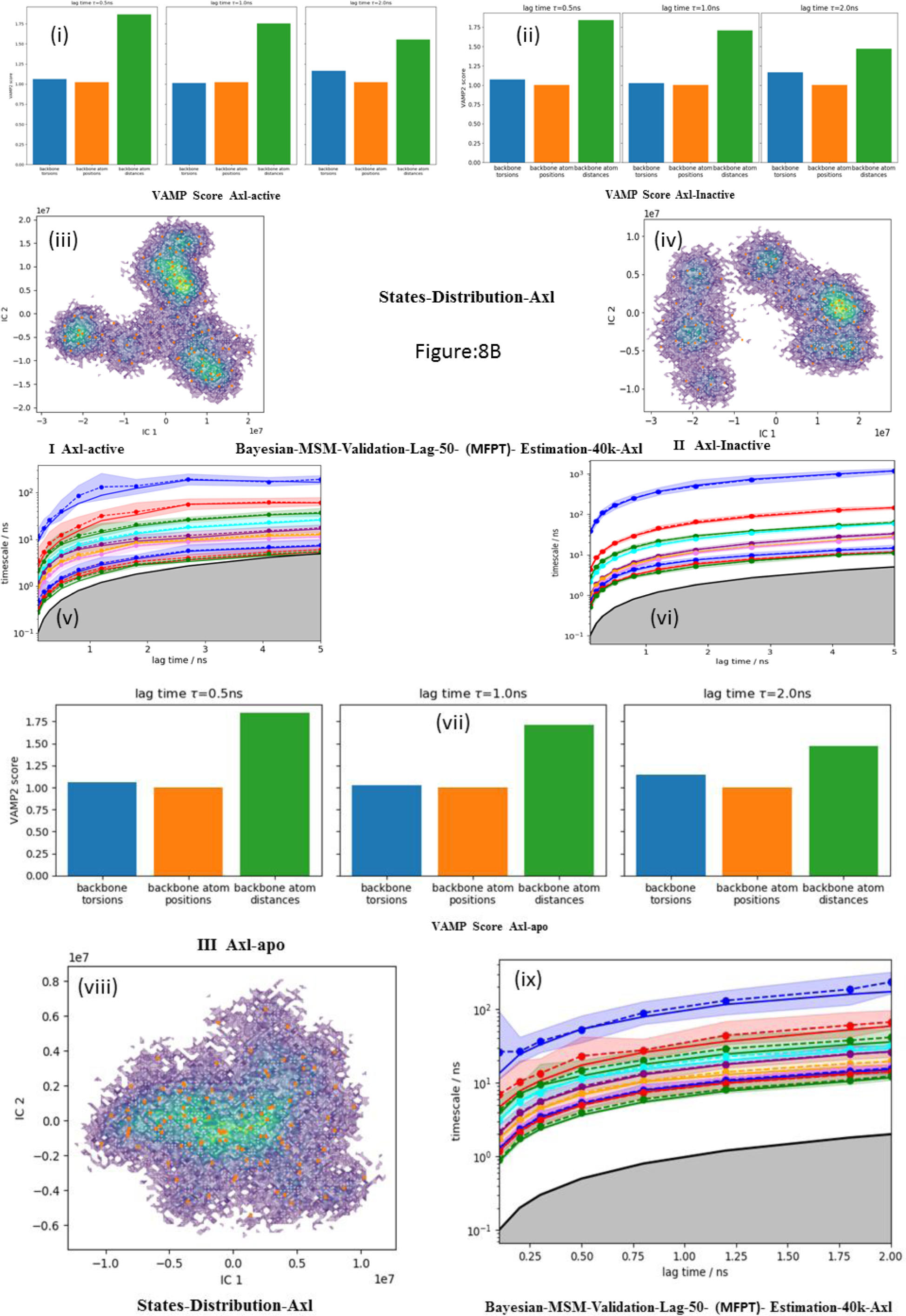
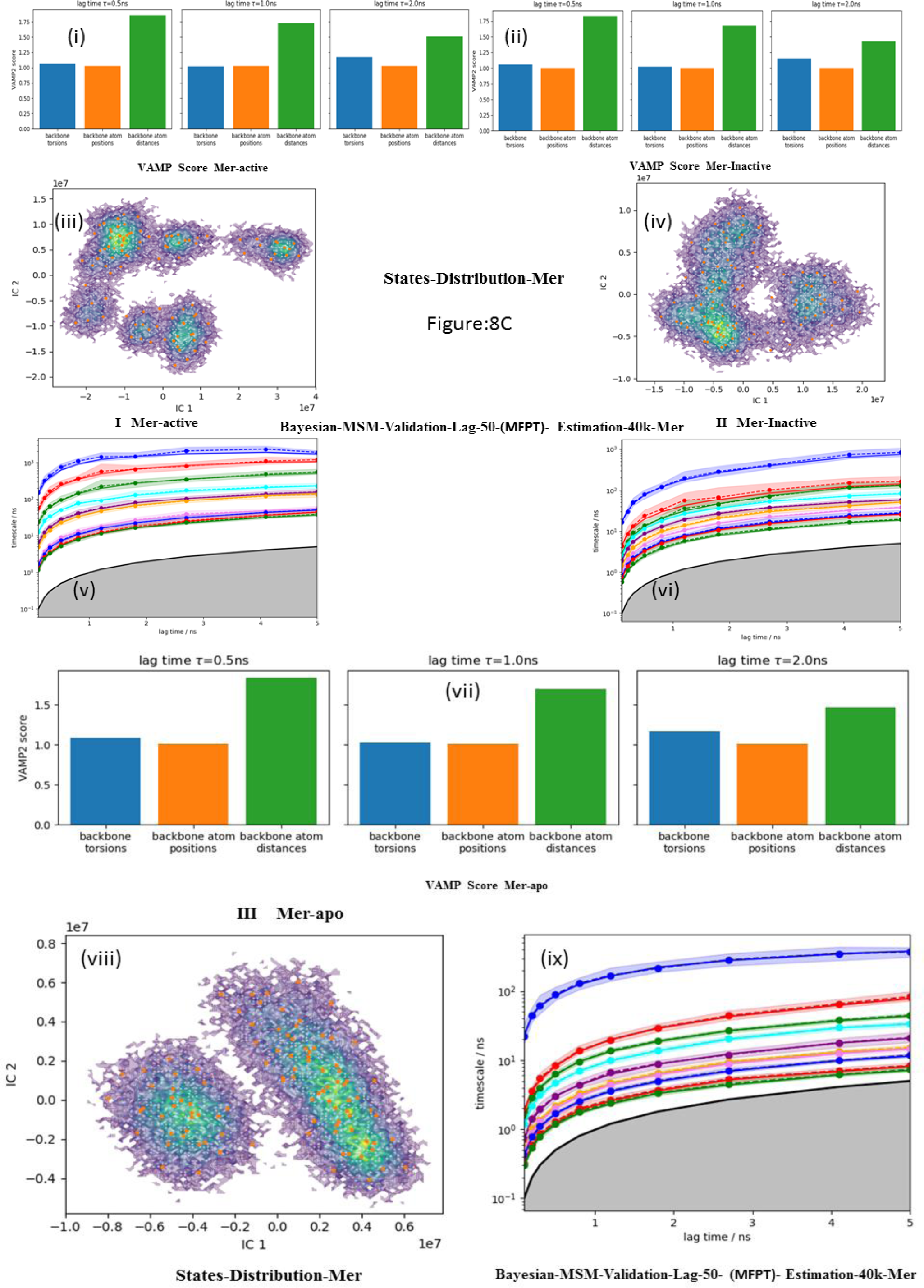
Sate Distribution analysis of TAM kinase apo & Cabozantinib bound active and inactive RTK from 1μs MD simulations. Markov state model validation [mean free passage time (MFPT) error bar analysis]. (8A) Tyro3 (8B) Axl (8C) Mer; (I) active; (II) inactive; (III) apo;

In the inhibitor bound form of TAM kinases, greater state distribution models co-exist in the active forms than in the inactive forms. The drug bound to kinase active state influences the kinetic signaling pathway more rather than the inactive state.^24^ Therefore, the active state kinase bound to inhibitor is more susceptible to arrest the dysregulated kinase activity (shown by the broken R-spine) in all kinetic HMM states. These observations provide key insights to describe that the kinase activity can be arrested through active state models of inhibitor bound RTK, where R-spine breaks in between activation loop and αC-helix in the active states. The hydrophobic surface R-spine is retained in the apo form of all the three TAM kinases. The fragmentation of R-spine occurred in a similar way in the cabozantinib bound Tyro3 RTK in active and inactive states, due to the increased distance between P-loop and αC-helix. This R-spine distortion in Tyro3 RTKs indirectly influences the number of active and inactive state distribution in equal proportions. In the inactive Axl and Mer RTKs, R-spine fragmentation also occurs between the P-loop and αC-helix, whereas in the active RTKs, R-spine fragmentation occurs between αC-helix and activation loop (Axl), and between P-loop, αC-helix and activation loop (Mer). These observations are shown in Figures 8 A to C. The discrete clustering of MSM estimation and validation was done with reversible estimation equilibrium transition probabilities. The discrete kinetic state models were further validated by analysis of hidden markov kinetic models. The implied relaxation timescales are extracted to validate the HMM in order to ensure the conditional transitions probabilities among 250 microstates. Therefore the implied timescale analyses indicated that the kinetic state distribution occurred within time intervals of a few nanoseconds range among 1 µs MD simulations time scale (**Figure S3A-C**).

The Mer active states have longer MD kinetic relaxation timescales among the active MSM kinetic forms of TAM RTKs. The inactive Axl kinetic state models have higher relaxation timescales within short range of time intervals. The critical observation from all TAM apo and inhibitor bound active and inactive kinetic states implied timescale plots, with 4.5 ns time scale separation is the average implied relaxation time scale among all. The Tyro3 apo has more relaxation time intervals than the rest of kinase systems (**Figure S3C**). The kinetic relaxation time intervals revealed that the inhibitor bound TAM RTK’s showed kinetic metastable state transitions due to various periodic time laps even though all TAM RTKs bound with same inhibitor (cabozantinib) (**Figure S3D**).

The MSMs of the members from same class of protein kinase complexes (TAM kinases bound to cabozantinib) is expressed as different relaxation time scale intervals obtained from the MD simulations. The free energy and stationary state distribution of apo Axl is higher than Tyro3 and Mer. From the table 1, we infer that there are unique kinetic Markov state models existing among them. These are classified as “kinetic non-equilibrium transition state models” (Tyro3 apo, Mer active, Axl inactive). This is further discussed in kinetic transition analysis. The lowest free energy and equal stationary distribution exist in stable kinetic model states of TAM kinases (Axl-active, Mer-inactive).

**Table 1:** Tyro3, Axl and Mer kinetic transition states analysis with specified free energy of nine HMM states. Red color indicates metastable kinetic equilibrium transitions states Blue color indicates metastable kinetic non-equilibrium transitions states

| Kinetic metastable states | Kinases Types |  | Axl |  | Mer |  | Tyro3 |  |
| --- | --- | --- | --- | --- | --- | --- | --- | --- |
| | | STATES | $\pi$ | G/kT | $\pi$ | G/kT | $\pi$ | G/kT |
|  | Active | 1 | 0.080859 | 2.515052 | 0.072653 | 2.622066 | 0.000000 | inf |
|  |  | 2 | 0.000000 | inf | 0.095574 | 2.347855 | 0.210675 | 1.557439 |
|  |  | 3 | 0.328240 | 1.114010 | 0.271887 | 1.302367 | 0.094495 | 2.359208 |
|  |  | 4 | 0.206730 | 1.576340 | 0.286168 | 1.251176 | 0.343637 | 1.068168 |
|  |  | 5 | 0.384171 | 0.956668 | 0.273718 | 1.295657 | 0.351193 | 1.046420 |
| Transition states |  |  |  | S4-S5 |  | S2-S4 |  | S4-S5 |
| | | STATES | $\pi$ | G/kT | $\pi$ | G/kT | $\pi$ | G/kT |
|  | Inactive | 1 | 0.128210 | 2.054082 | 0.032751 | 3.418814 | 0.030226 | 3.499054 |
|  |  | 2 | 0.262320 | 1.338189 | 0.037707 | 3.277907 | 0.058256 | 2.842902 |
|  |  | 3 | 0.138915 | 1.973891 | 0.260523 | 1.345064 | 0.132997 | 2.017429 |
|  |  | 4 | 0.171230 | 1.764746 | 0.151968 | 1.884087 | 0.255189 | 1.365750 |
|  |  | 5 | 0.299324 | 1.206230 | 0.517051 | 0.659614 | 0.523331 | 0.647540 |
| Transition states |  |  |  | S2-S5 |  | S4-S5 |  | S4-S5 |
| | | STATES | $\pi$ | G/kT | $\pi$ | G/kT | $\pi$ | G/kT |
|  | Apo | 1 | 0.008222 | 4.800930 | 0.081258 | 2.510125 | 0.078556 | 2.543949 |
|  |  | 2 | 0.052043 | 2.955685 | 0.239732 | 1.428232 | 0.083054 | 2.488261 |
|  |  | 3 | 0.080056 | 2.525032 | 0.131020 | 2.032406 | 0.139106 | 1.972522 |
|  |  | 4 | 0.165858 | 1.796625 | 0.228406 | 1.476629 | 0.293660 | 1.225334 |
|  |  | 5 | 0.693821 | 0.365541 | 0.319583 | 1.140738 | 0.405625 | 0.902326 |
| Transition states |  |  |  | S4-S5 |  | S4-S5 |  | S3-S5 |

The kinetic transition state between Axl active and Mer inactive has higher free energy and approximately equal stationary distribution values (Tyro3 active/inactive) and are classified as “kinetic equilibrium transition state models”. As per the state distribution difference between active-inactive states of Axl inactive HMM has half (1/2) of the stationary distribution of Axl active (more active state distribution). The inactive Mer has ¾ of the state distribution of active Mer RTK. The Tyro3 has equal contribution in active and inactive stationary distributions among kinetic HMM states. The surface free energy of Axl has same energy values in active and inactive states (∼ 4.0 kcal/kT per 5 states-Axl) but Tyro3 and Mer have 0.5 kcal and 1.2 kcal, respectively per five MSM states energy difference between the active and inactive hidden Markov states (**Figure 9A-C**). Each hidden MSM state contains five metastable kinetic conformers from sampling of 40K conformers to study MSM validation.

**Figure 9:**
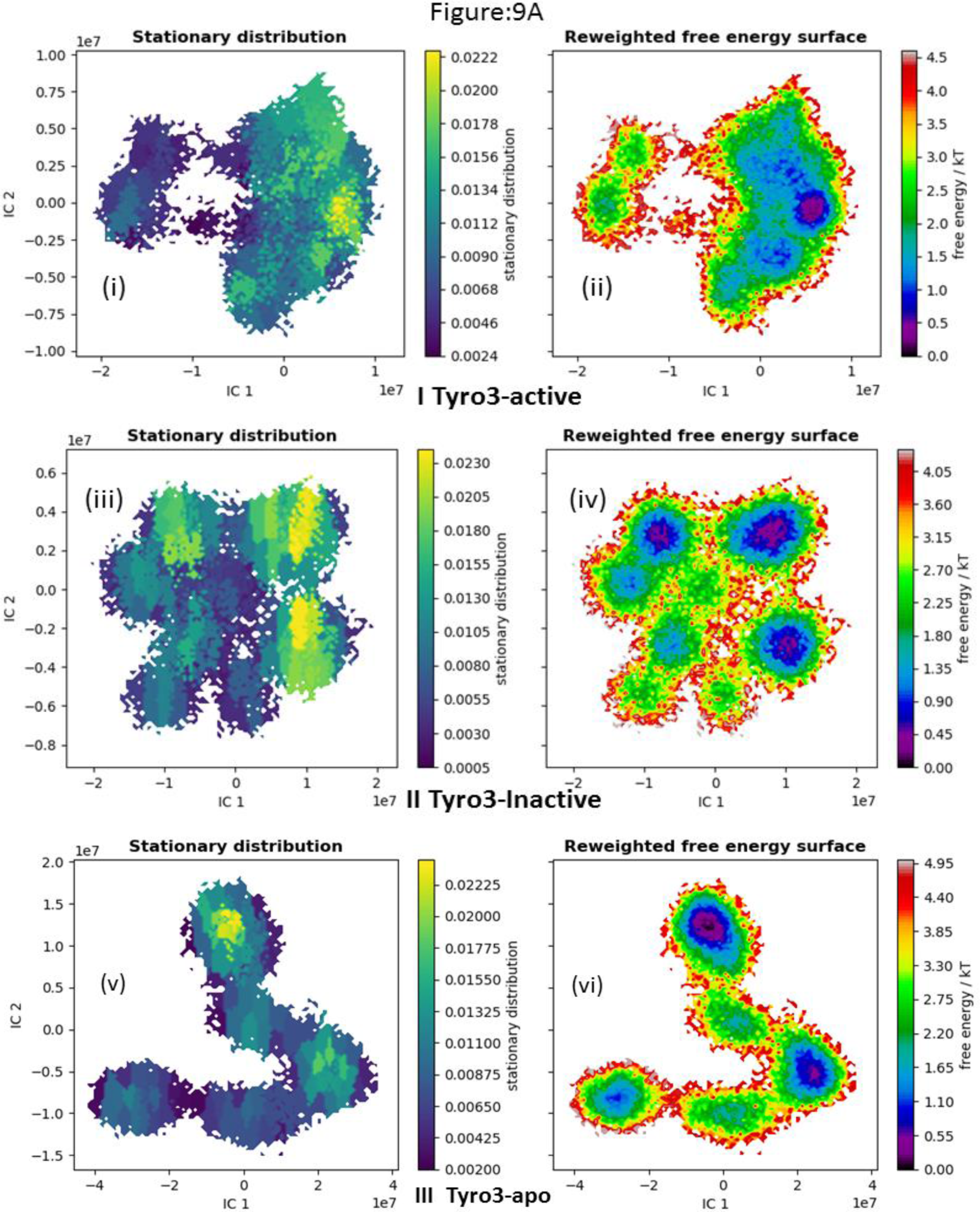
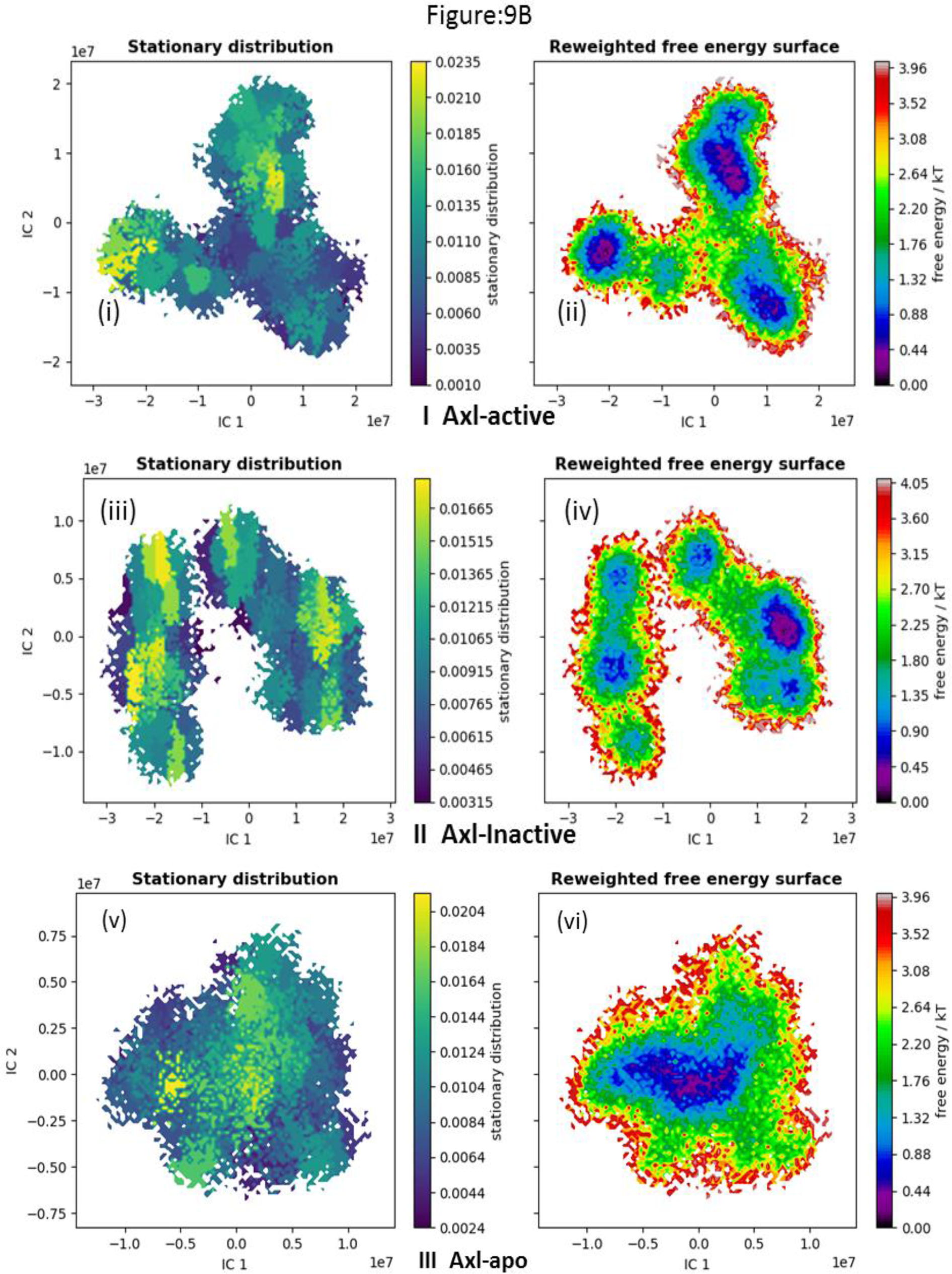
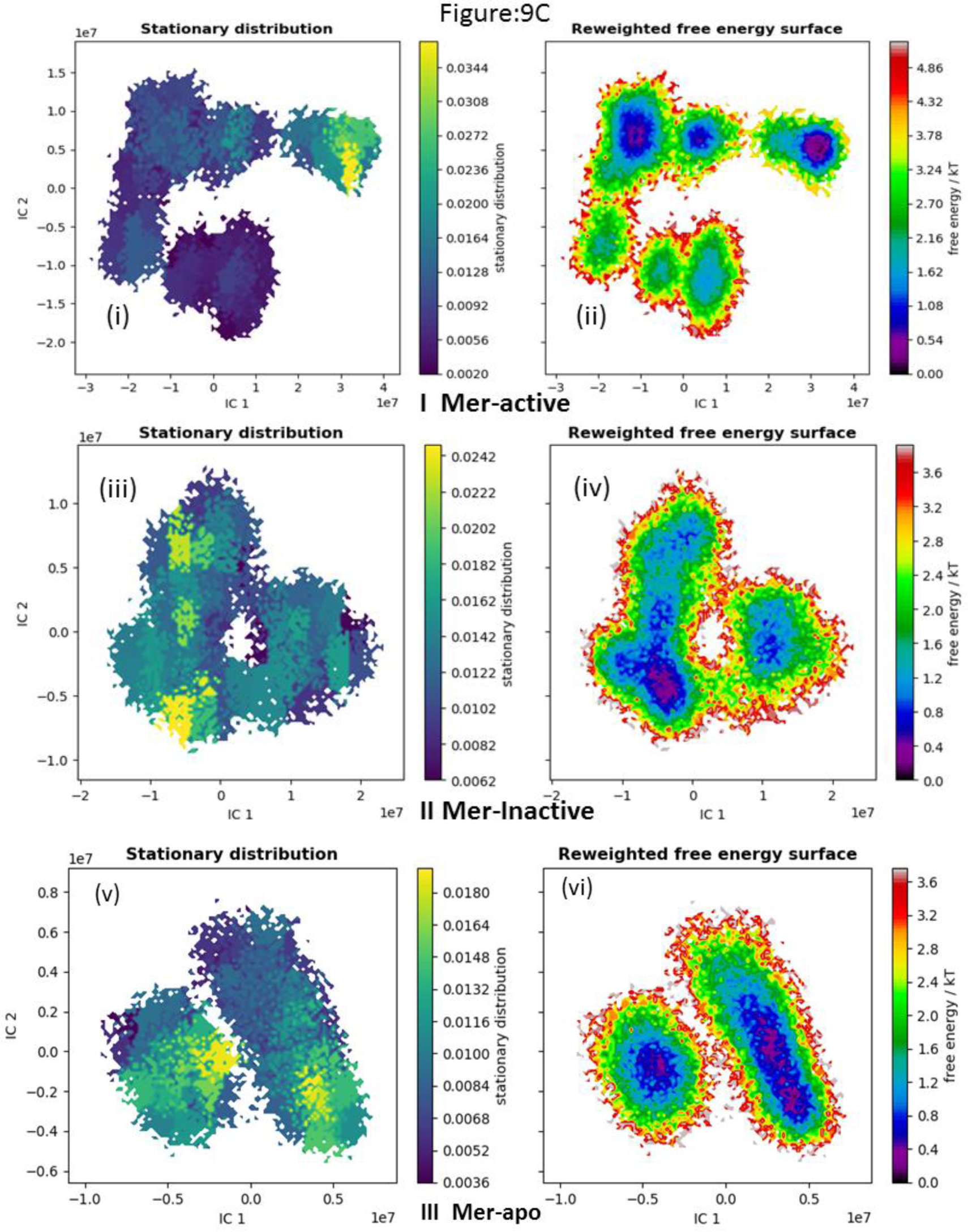
Stationary distribution and Free energy surface analysis of TAM kinase apo & Cabozantinib bound active and inactive states of RTK from 1μs MD Simulations. (9A) Tyro3 (9B) Axl (9C) Mer (I) active; (II) inactive; (III) apo;

### Kinetic transition state analysis

The estimated five state kinetic metastable models were designed based upon active space distribution of HMMs of TAM RTKs. All the five kinetic metastable state transitions occurred based upon kinetic transition energy (Table-1). The apo Axl has higher transition energy (4.8 kcal), inactive Mer (3.4 kcal) and inactive Tyro3 (3.5 kcal). Out of the nine kinetic states, six kinetic transition states are represented as metastable kinetic equilibrium transition states as these kinetic transitions occurred in **S_4_-S_5_** states. The metastable kinetic non-equilibrium transitions exist in various types of kinetic metastable states (Tyro3 apo – **S_3_-S_5_**; Mer-active- **S_2_-S_5_**; Axl inactive- **S_2_-S_4_)** from the nine metastable transition states (**Figure 10A-D**). All non-equilibrium kinetic transitions occur with a very low transition energy (2 - 2.6 kcal). All these hidden states are classified based upon kinetic transition energy and state transitions. All the metastable kinetic equilibrium transitions occurred with a high energy (2.3-4.8 kcal) (**Figure 11A-F**). As per the individual RTK, the Axl apo kinase has higher kinetic transition energy among all TAM RTKs in apo and inhibitor bound active and inactive forms. The next higher kinetic transition energy exists for Mer and Tyro3 inactive forms. It is clearly evident that all inhibitor bound RTKs exhibit different kinetic metastable states in the overexpressed RTKs during the protein function. According to approximate difference in transition probability of active to inactive metastable kinetic states in Tyro3, Axl, and Mer RTKs, for Tyro3, 1^st^ MSM state has higher transition probability difference (50 %), for Axl and Mer RTKs, 2^nd^ MSM states have higher transition probability difference (**Figure 11G-I**).

**Figure 10:**
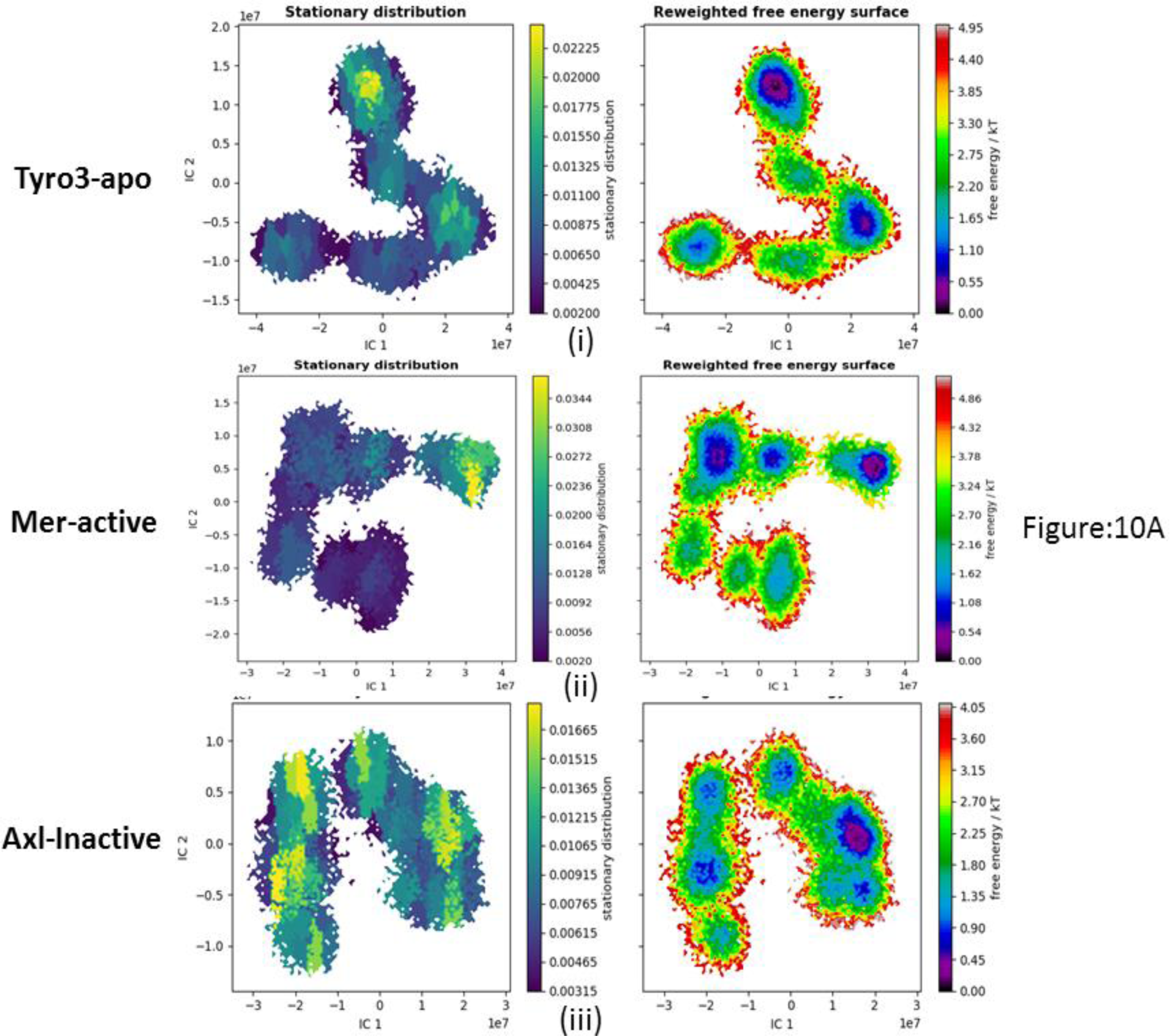
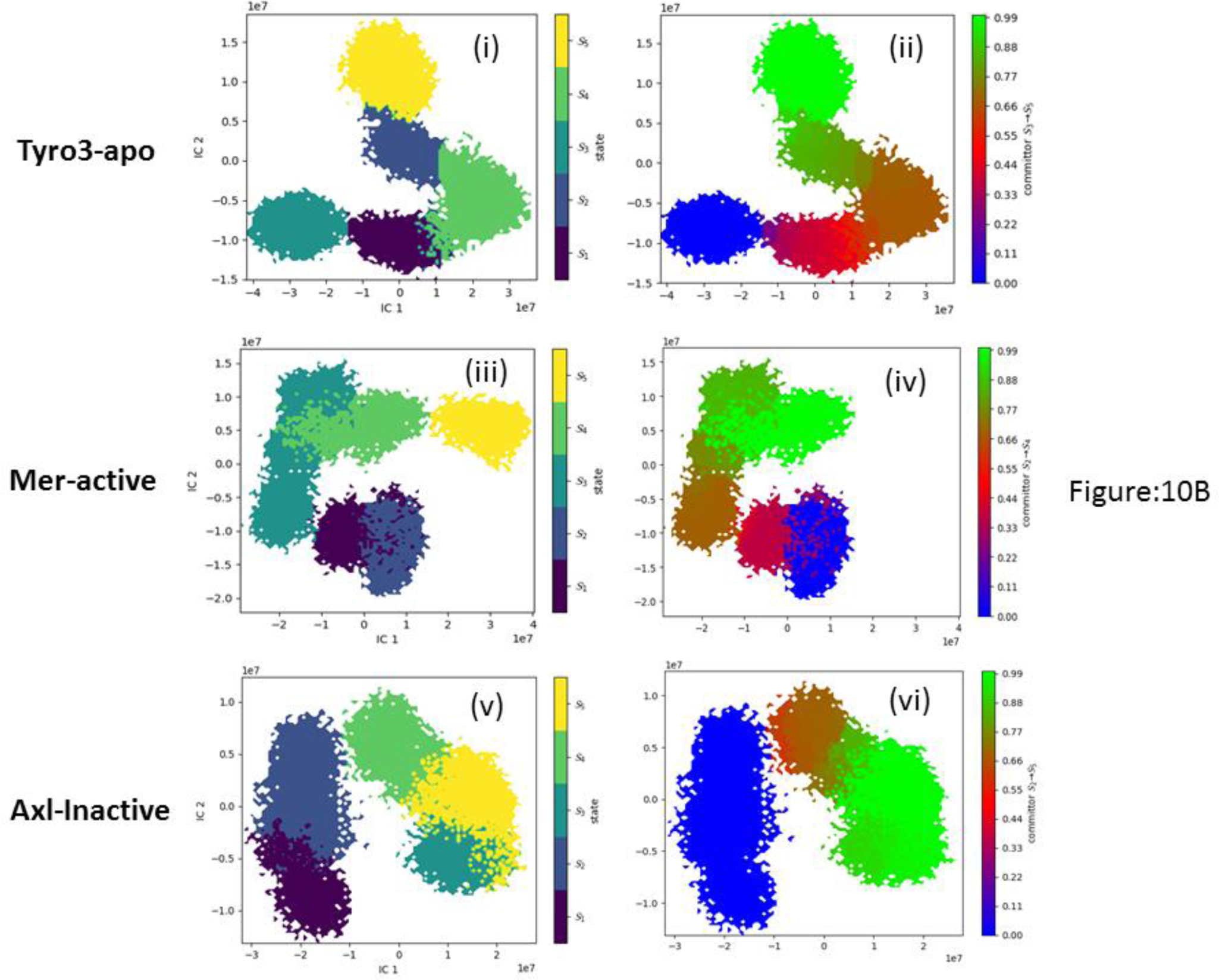
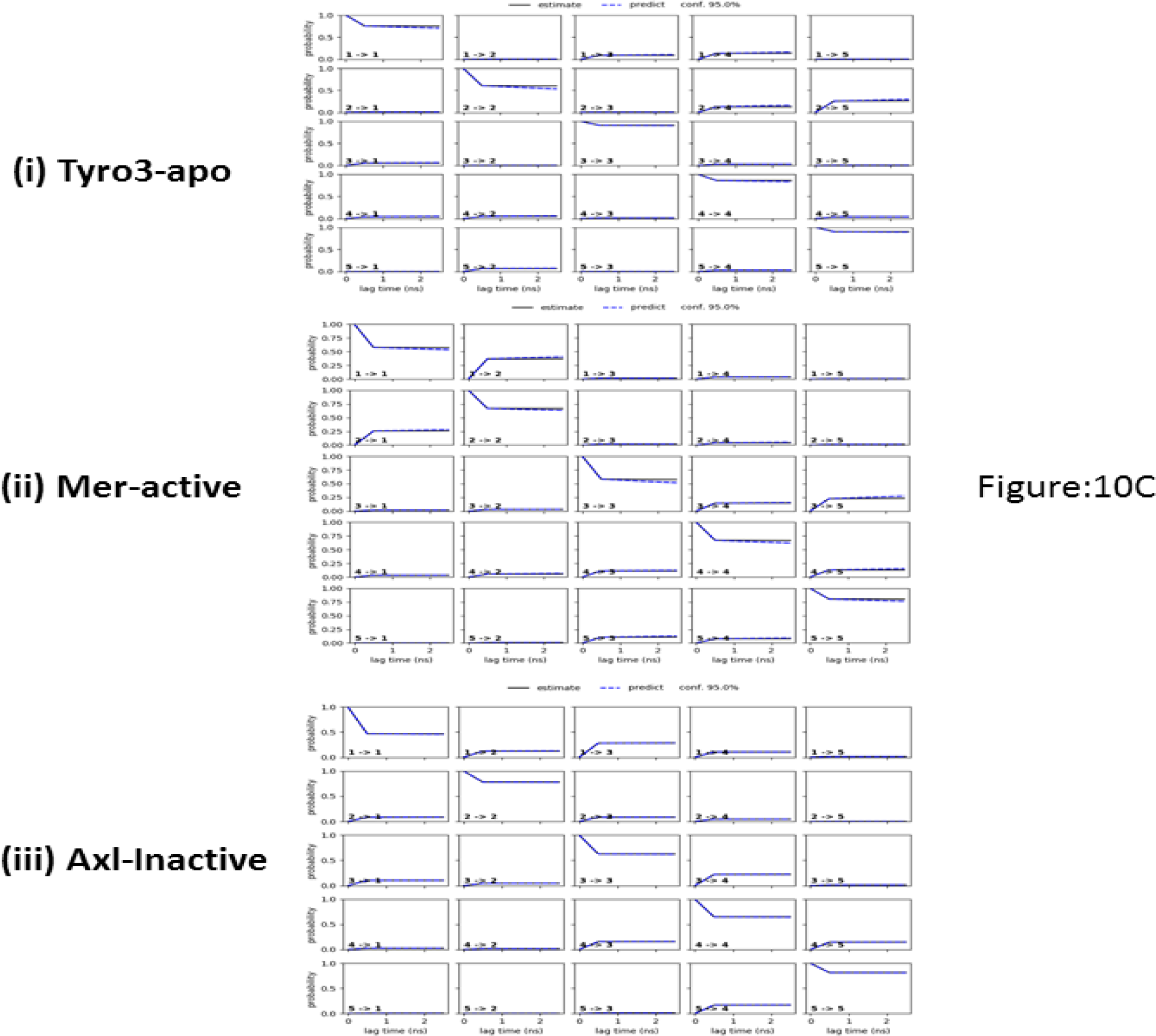
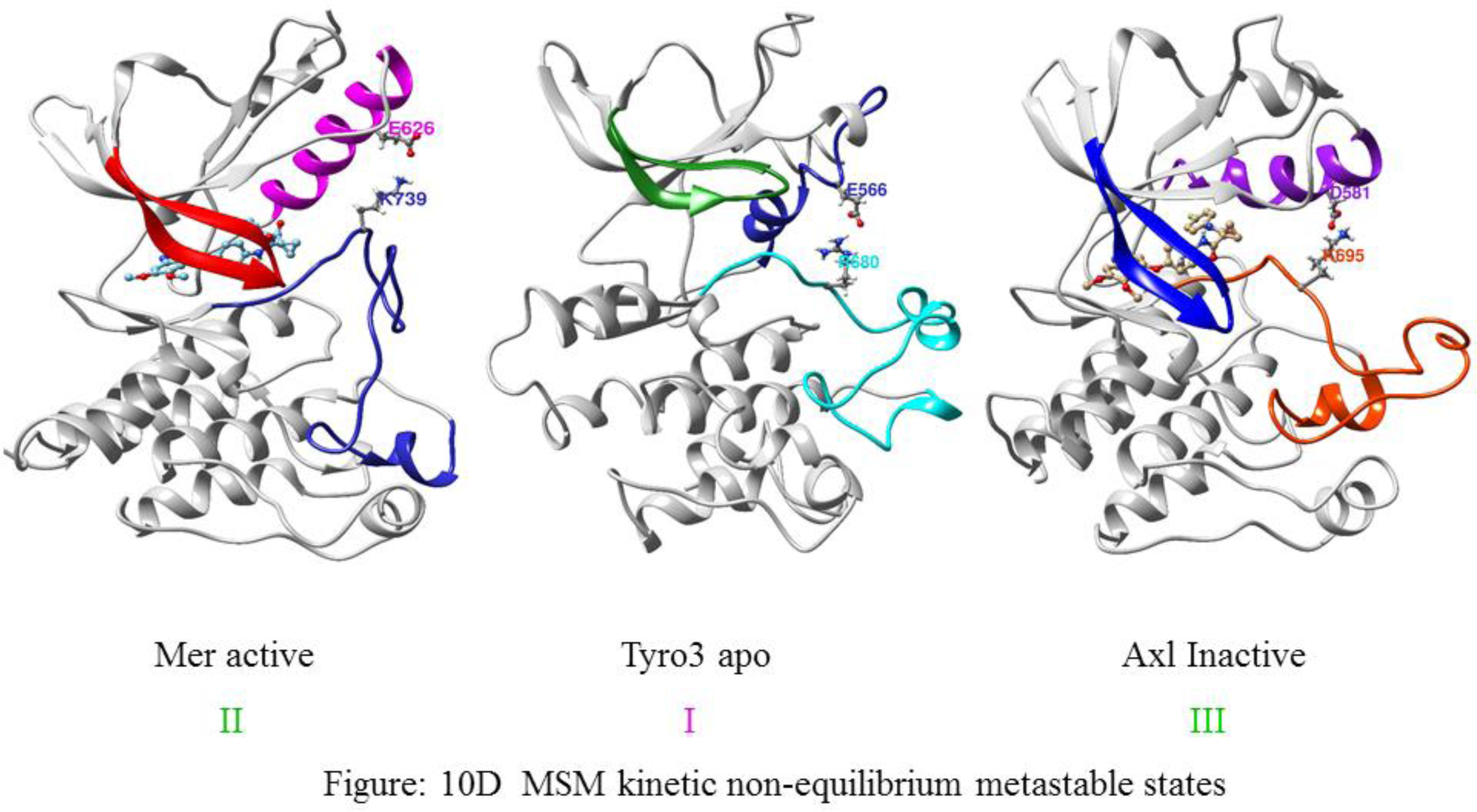
Kinetic metastable five states estimation and Kinetic metastable transition state analysis of metastable kinetic non-equilibrium transitions states Tyro3-apo; Mer-active; Axl-inactive Tyro3 apo and Cabozantinib bound Mer-active and Axl-inactive states of RTK from 1μs MD simulations. (10A) Stationary states and Reweighed free surface energy of non- equilibrium transitions states (10B-C) MSM five states estimation & Kinetic transition states & their transition probabilities (10D) Specific states distance between side chains of Asp/Glu-αC-helix – Lys/Arg-Activation-Loop pairs in Tyro3 apo and Cabozantinib bound Mer-active and Axl-inactive

**Figure 11:**
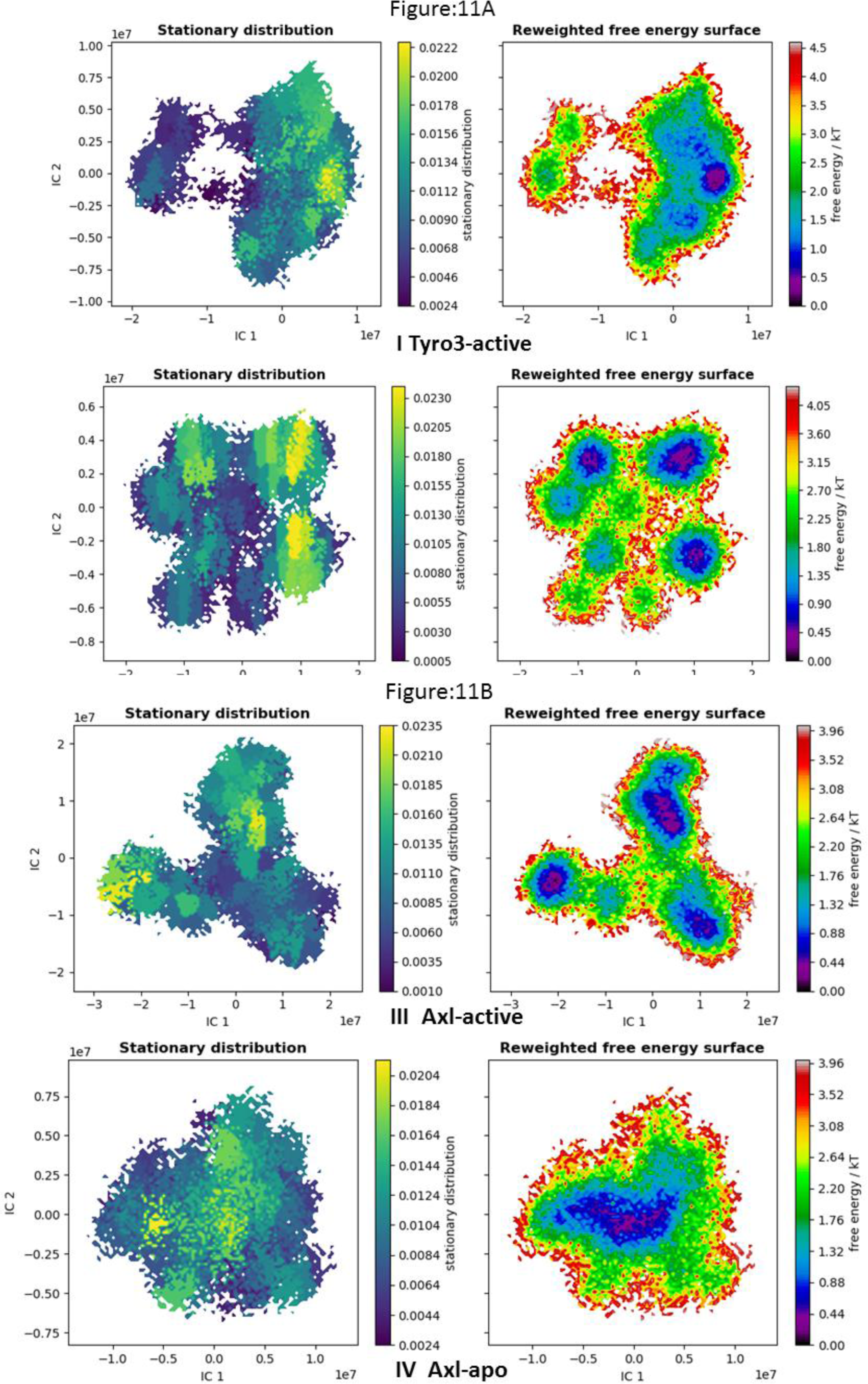
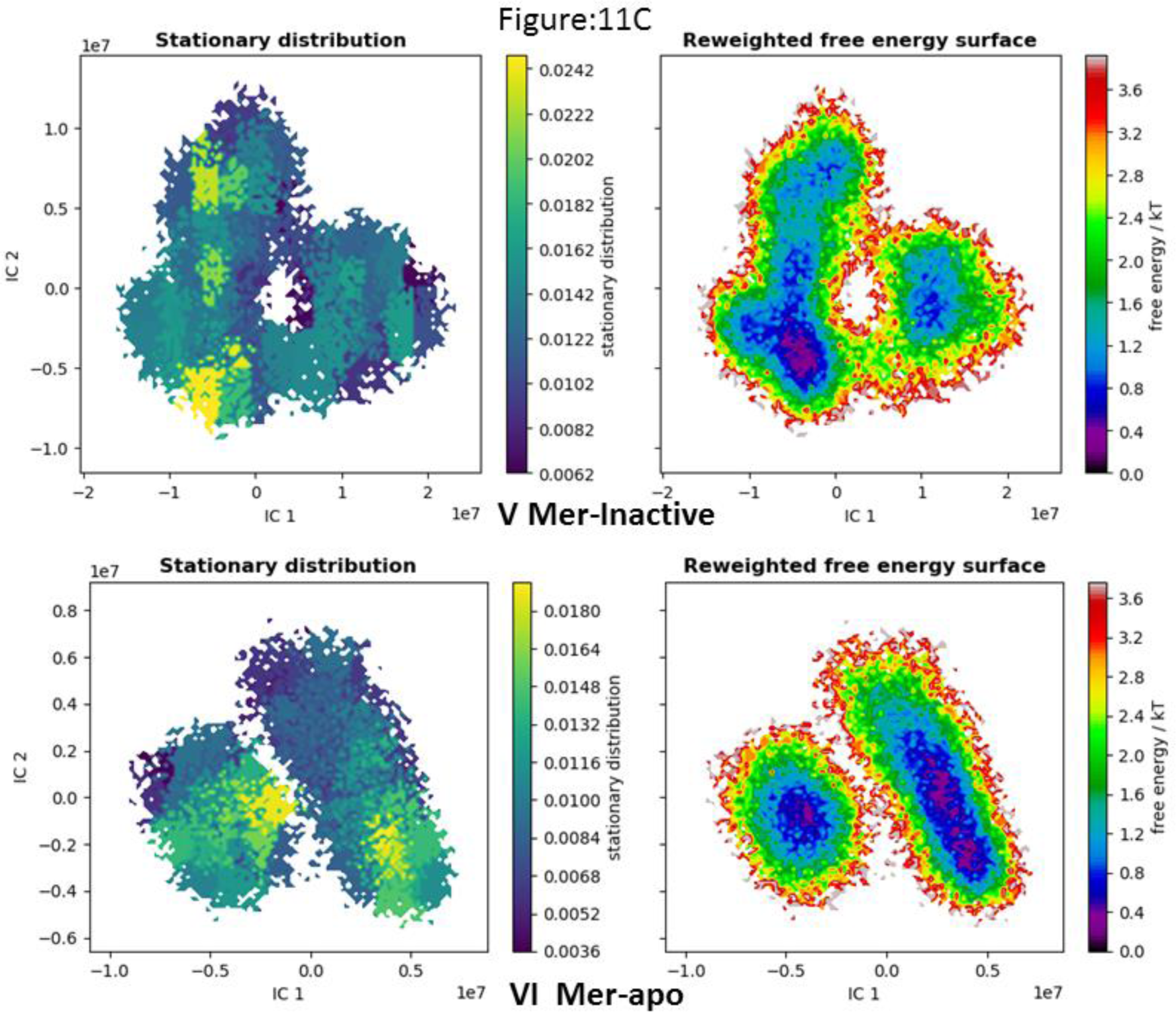
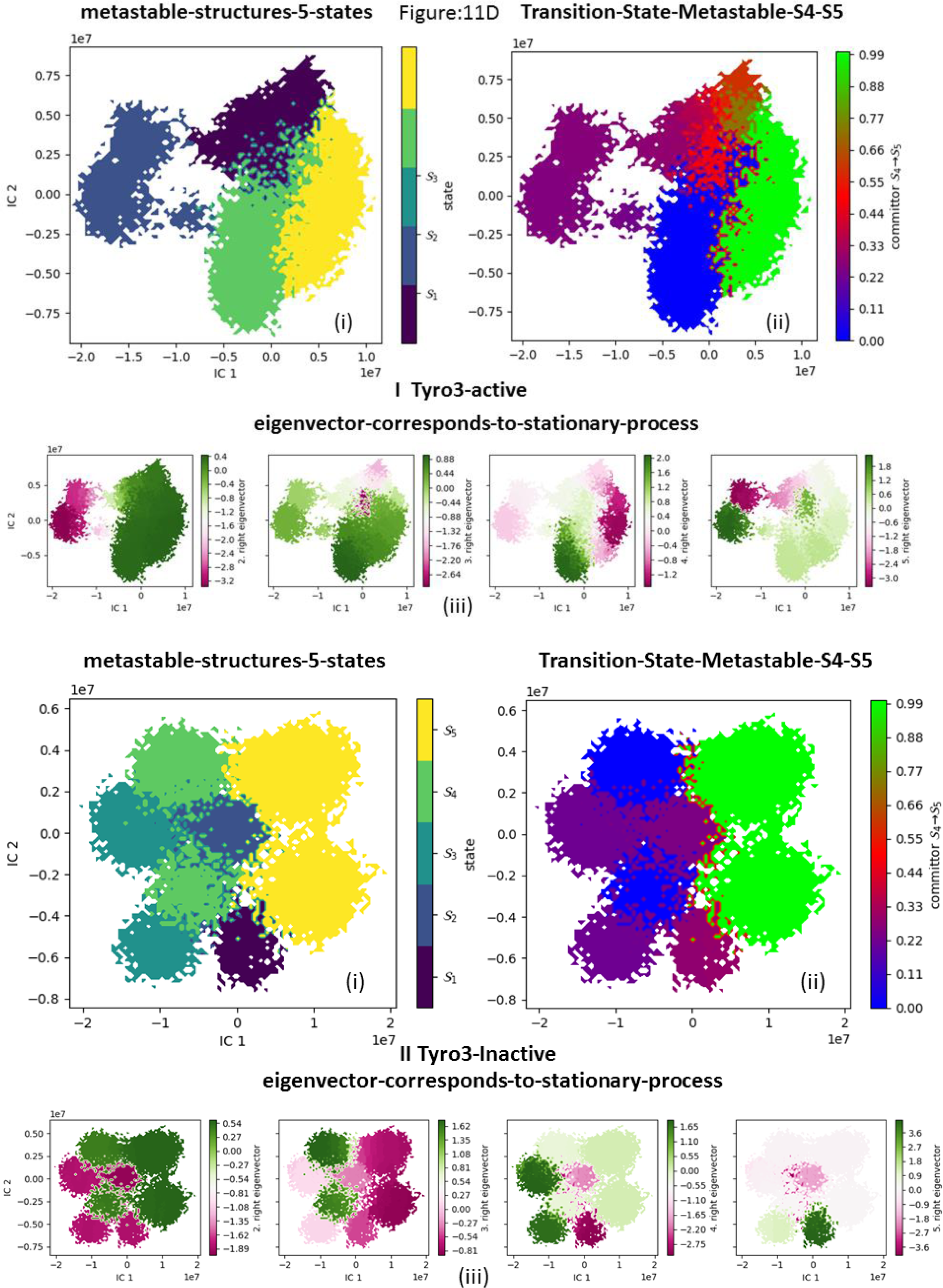
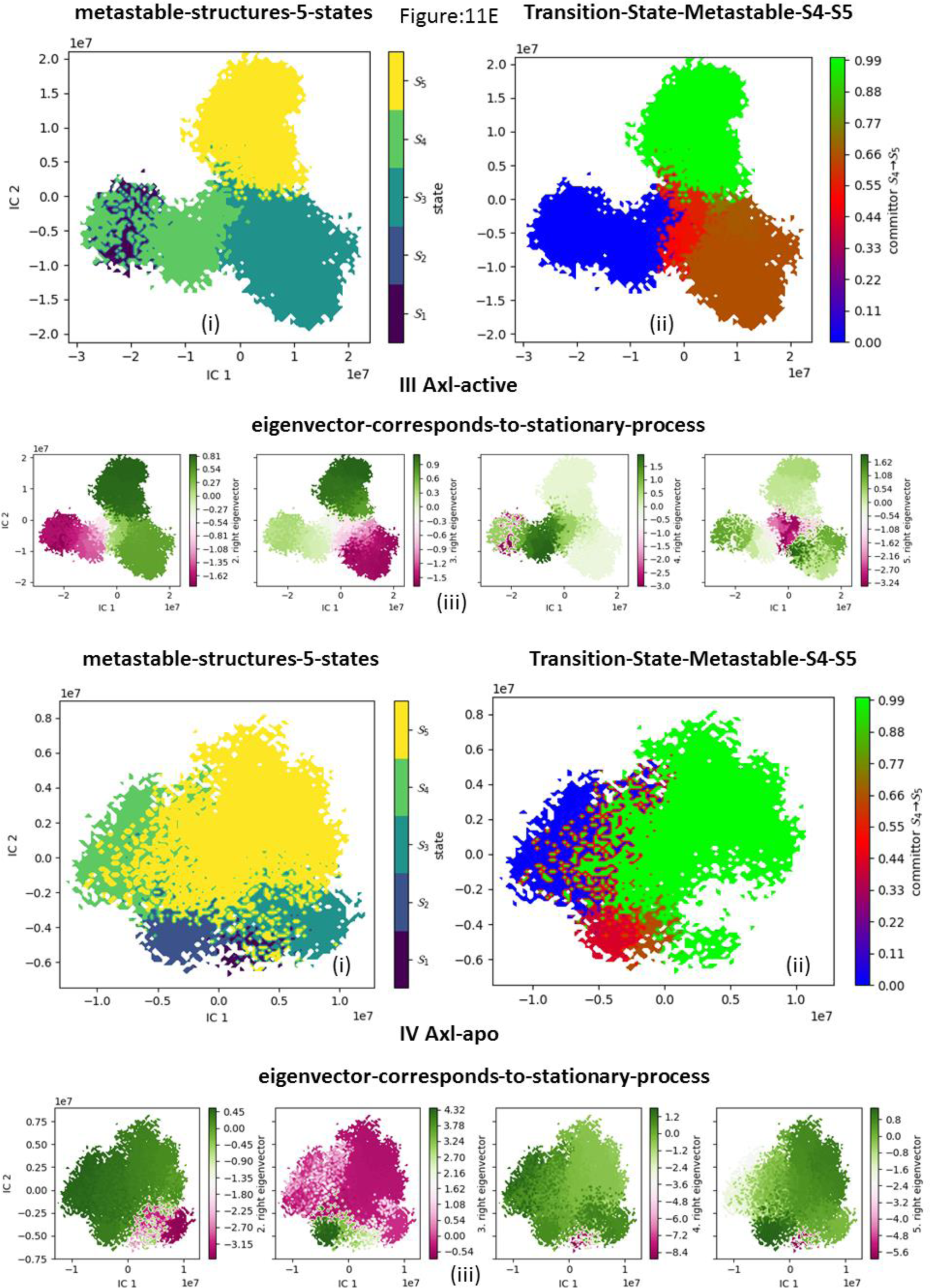
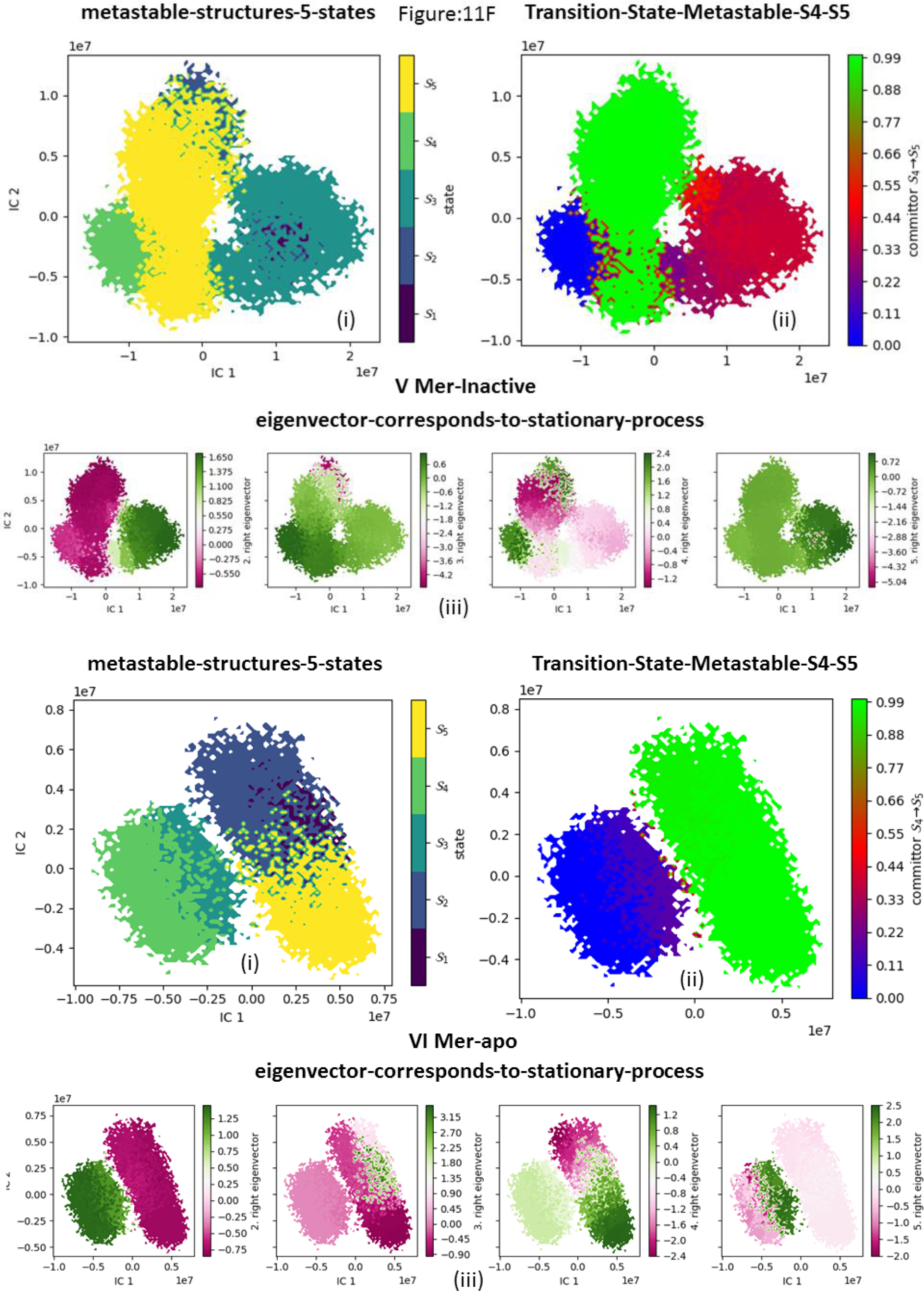
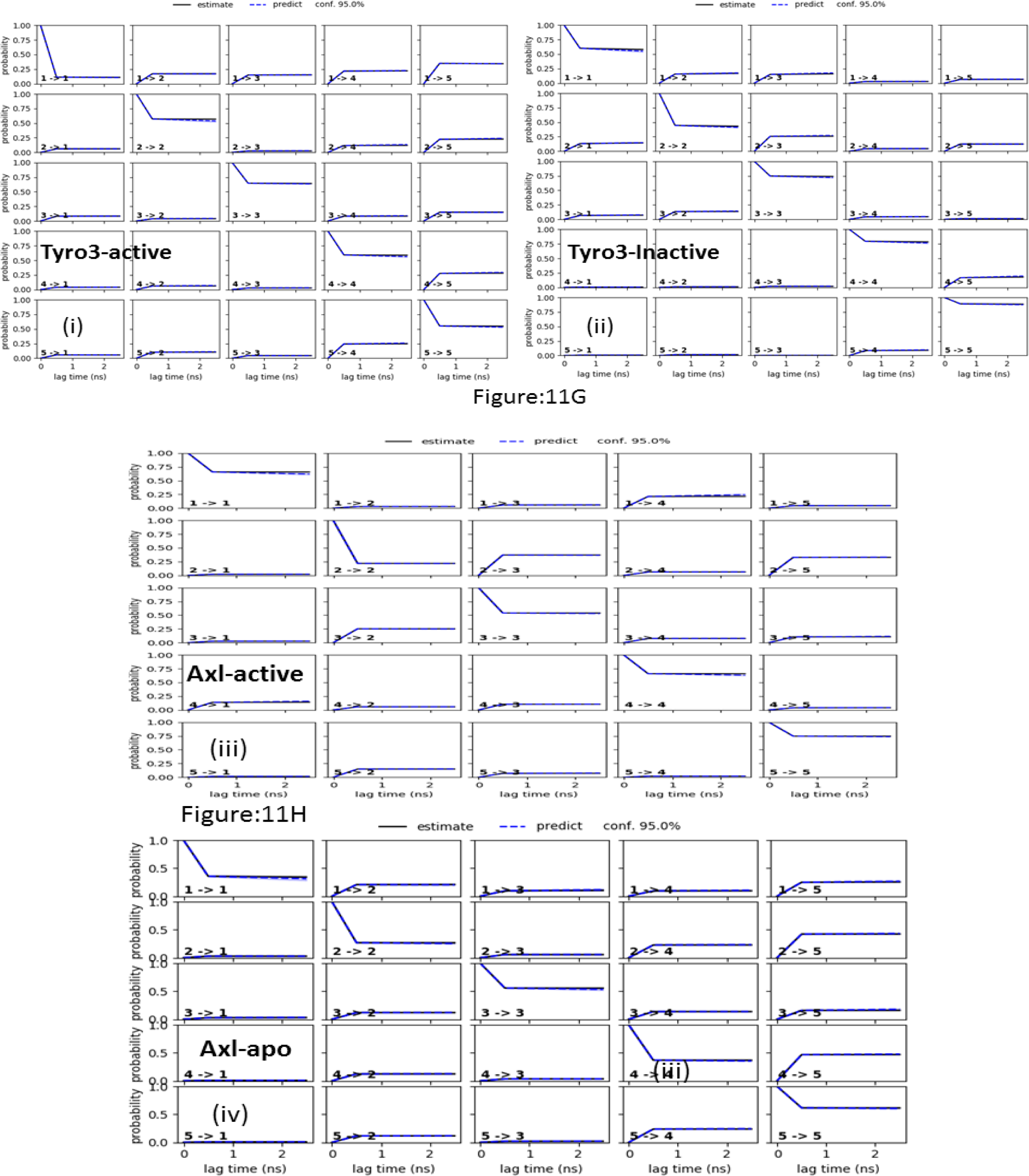
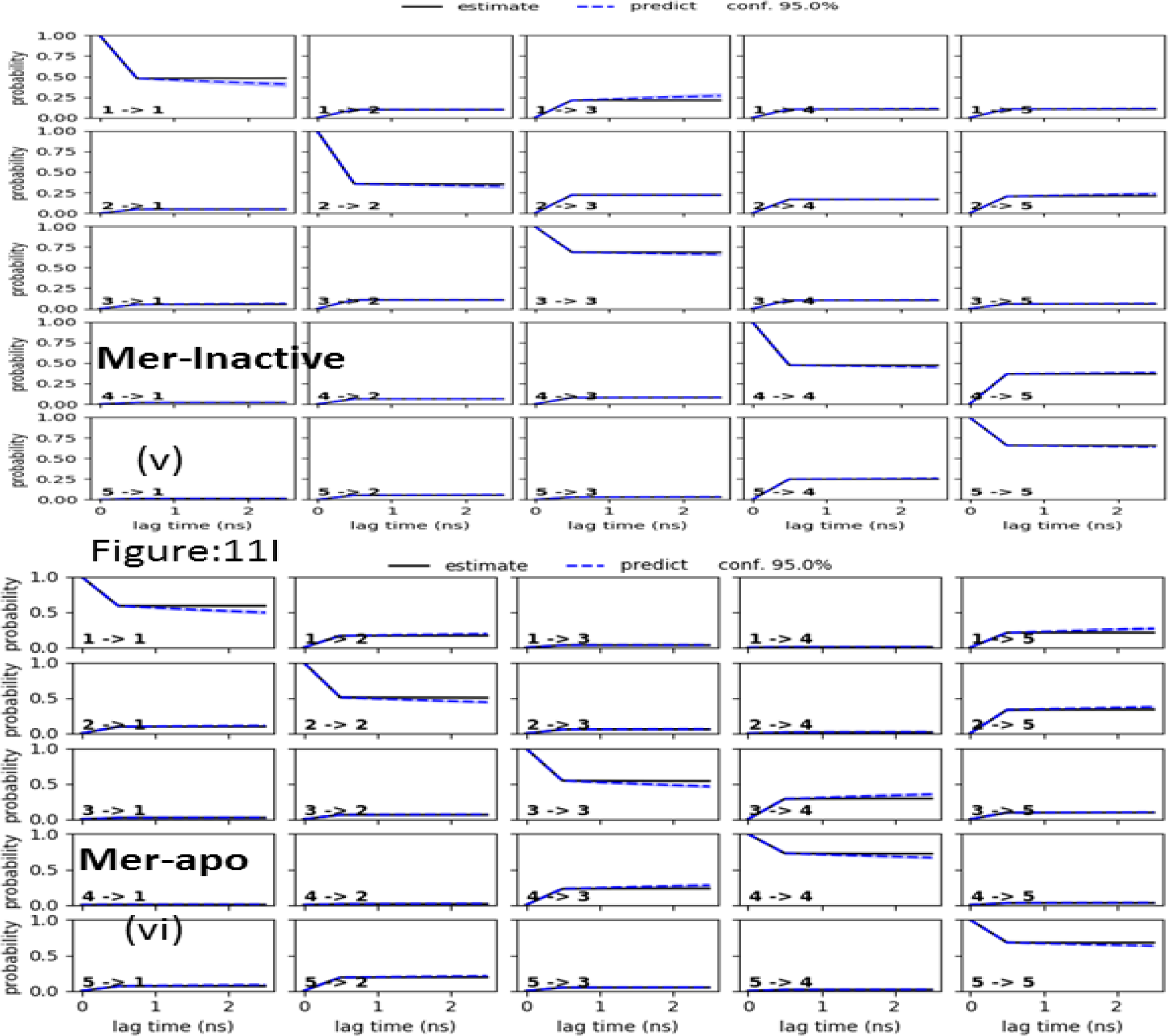
Kinetic metastable five states estimation and Kinetic metastable transition State analysis of metastable kinetic equilibrium transitions states of RTK from 1μs MD simulations. (I) Tyro3 -active; (II) Tyro3-inactive; (III) Axl-active; (IV) Axl-apo; (V) Mer-inactive; (VI) Mer-apo;. (11A-C) Stationary states and Reweighed free surface energy of equilibrium Transitions states. (11D-I) MSM five states estimation & Kinetic transition states & their Transition probabilities.

**Figure 12:**
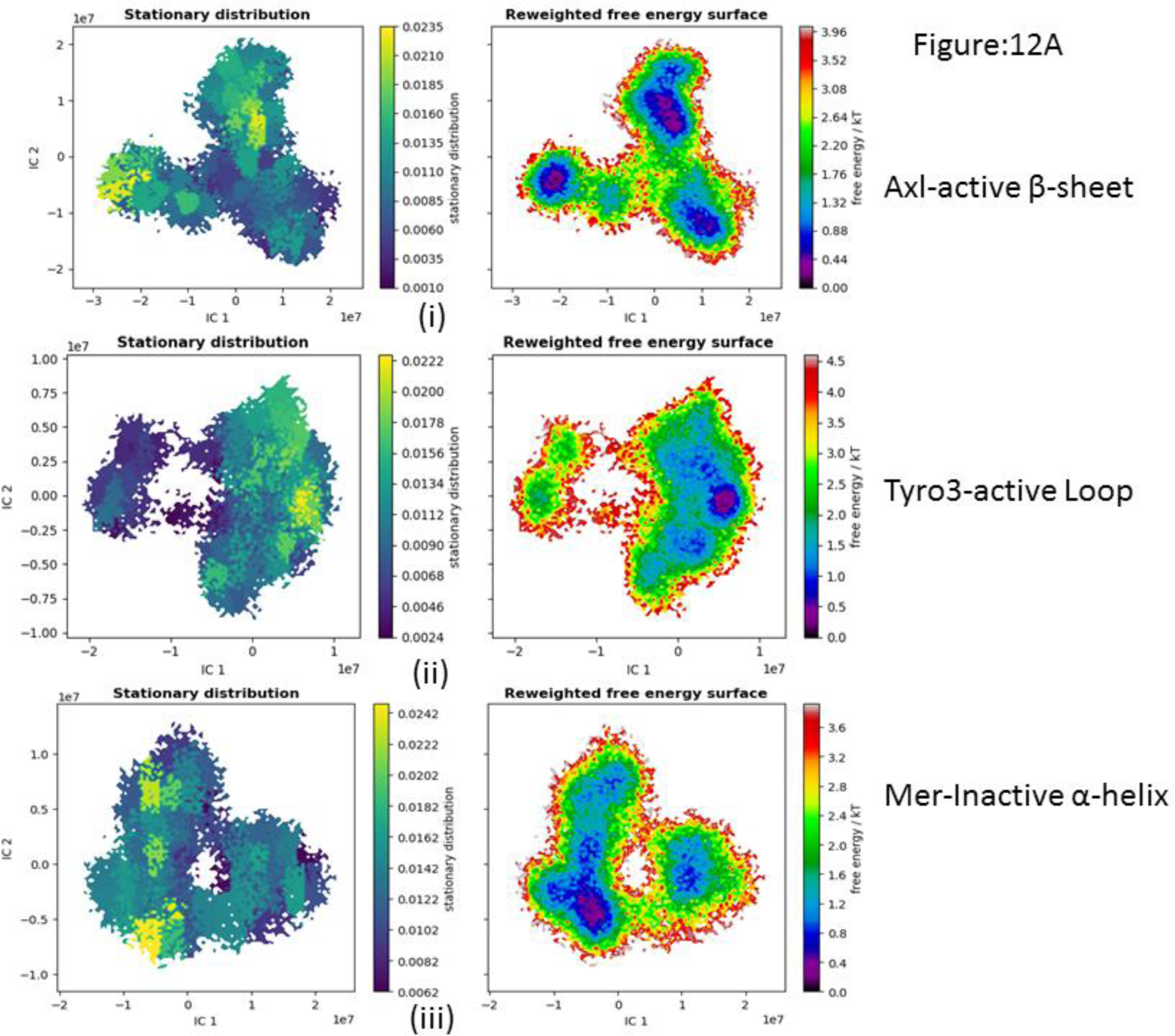
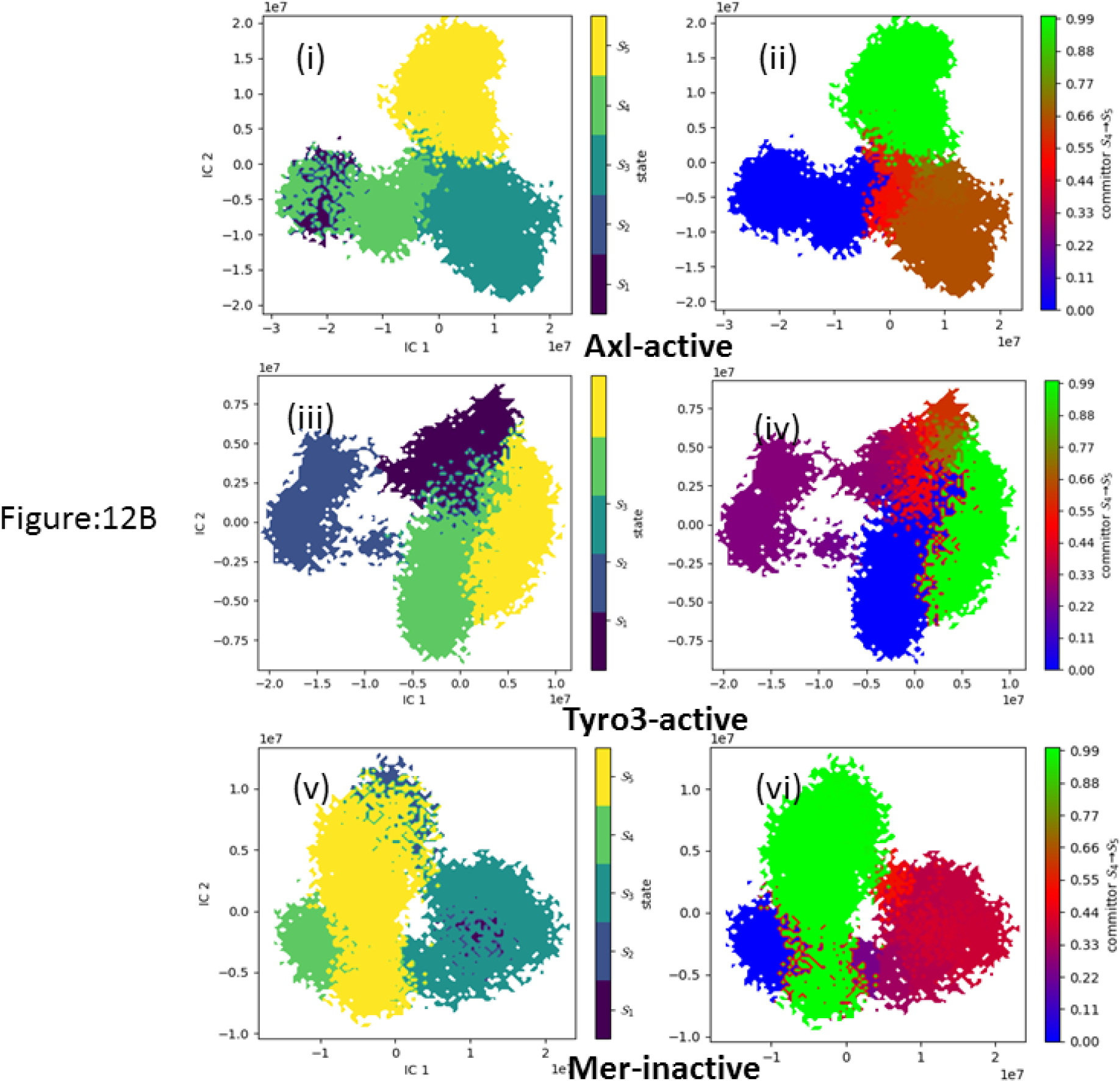
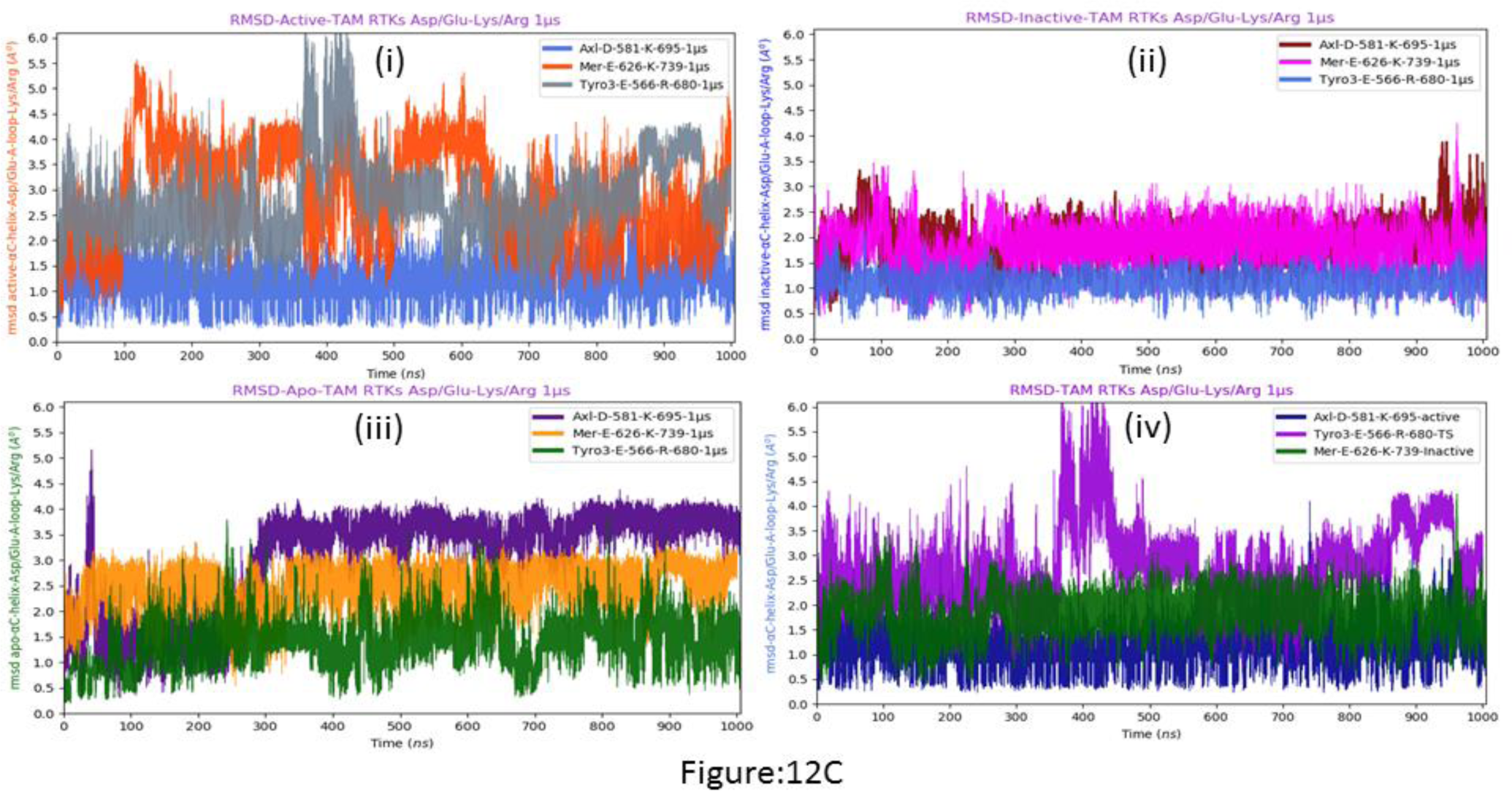
The selective kinetic equilibrium transitionstate models among nine Hidden markov state models of TAM RTKs. (12A) Stationary distribution and Free energy surface analysis of three kineticequilibriumTransition state models. (i) Axl-active; (ii) Tyro3-active; (iii) Mer-inactive; (12B) Kinetic metastable five states estimation and Kinetic metastable transition stateAnalysis three kinetic equilibrium transition state pairs. [All kinetic TS S_4_-S_5_] (12C) Distance plots between Asp/Glu-αC-helix – Lys/Arg-Activation-Loop in 3 kineticEquilibrium transition state pairs of Cabozantinib bound active and inactive states Of RTK from 1μs MD simulations. (i) Active; (ii) Inactive; (iii) Apo; (iv) TAM

The transition of active kinase state to inactive state can be explained based upon kinetic metastable states of these specified MSM conformer analysis.^25^ From the figure it can be seen that the stationary state distributions in Axl active is double when compared Tyro3 active, and Mer inactive states has only 4/3 proportion. Therefore, Axl active RTK has more active stationary states. The relative transition state probability is explained on the basis of salient feature analysis in hidden Markov kinetic states. These could be key intermediate structures among subfamily of TAM RTKs. However, these protein kinases driven from active to inactive states expressed significant structural changes upon binding with inhibitor. The active state of Axl kinase consists of activation loop that transits from β sheet to helical structure in the inactive state. Mer RTK shows high structural changes in activation loop which converts from loop (active) to helical (inactive) in their respective state transitions (2-2 transition probability), while Tyro3 does not have any significant change in MSM kinetic states.

## Discussions

### Mechanistic strategy of TAM RTKs complexed with cabozantinib

RTK domain can switch from active to inactive states and vice-versa due to either inhibitor binding, or influences of the R-spine or C-spine during MD simulations at longer timescales. We observed salient changes in the spatial conformational states due to inhibitor binding to the active site during MD simulations in various regions of Tyro3, Axl and Mer kinases. These specified regions of spatial orientations are not directly observable from the conventional RMSD plots.

The inhibitor bound Axl kinase activation takes place in the transition of active-inactive kinetic models. At the specified time scales of MD simulations, these states coexist with broken R-spine in the active and inactive metastable states. In addition, the RMSD of Axl differentiates due to the coexistence of active-inactive states throughout 1 µs time scales (**Figure 2a**). The RMSD of specific loops in Axl is observed at higher square fluctuations occurring at the loop connecting β4-β5 strands (Glu609-Pro614), αC-helix and activation loop (**Figure 4d, 4e, 4f**). It is clearly evident from the RMSF plots that cabozantinib drug binding influences the inactive state of Axl and Mer kinase more than their active states (**Figure 5**). When cabozantinib binds the kinetically metastable states of TAM RTKs, it arrests the mechanism of kinase activity by inhibiting the up-regulation of its enzymatic activity.

Based upon individual RMSD plots of the R-spine and activation loop (**Figure 4b, 4c**), it is clear that cabozantinib binding influences the activation loop and hydrophobic spine in individual kinetic states. A specific spatial conformational variation in RTK’s occurs only in the activation loop and R- spine. The active and inactive forms of apo conformers of TAM kinases appeared to have intact R-spine. This could lead to the normal signal transduction process while it is attached the regular ligands [GAS-6 and Pros1] bound to the extracellular regions of TAM RTKs. The R-spine of TAM apo form compared with 1 µs trajectory MD conformers of TAM-inhibitor complex have specific hydrophobic surface broken in either of active and inactive kinetic states (**Figure 2 a-c**).

The kinetic transition probabilities of 40K frames of dynamic kinetic metastable states for each of the protein complex trajectory was obtained from AMBER MD simulations data. The five metastable states from Chapman-Kolmogorov test described transition probability from one kinetic metastable state to another with 95% confidence (**Figure 11G-I**). Combining all these transitions probabilities with transition states and assigning five sampled metastable states could provide good insights and predict long lived transition states in MD simulations trajectories with perron-cluster cluster analysis (PCCA++) clustering algorithm.^26–28^

The RTKs are involved in signal transduction process in which dysregulated kinase protein is inhibited such that the cells initiate programmed cell death with help other kinases and proteases belonging to the caspase enzyme.^29^ The regulated and dysregulated kinase can be distinguished with help of R-spine **(Leu-620 {β4-sheet P-loop}; Met-589 {αC-helix}; Phe-691 {DFG-A-loop}; His-670 {Catalytic loop})** closed and broken conformers of apo and inhibitor bound TAM RTKs, respectively due to significant conformational changes. Regulated kinase has closed and continuous R-spine in both active apo form in RTKs.^30^ But inhibitor bound dysregulated kinase experiences large conformational deformations in their regular structures due to the influence in certain parts of RTKs with overwhelmed hidden dynamic states to trigger kinase domain equilibration between active and inactive states. Indeed, the drug (cabozantinib) bound at RTKs active site, triggers the activation loop folding into either β-sheet (active-state) or α-helix (inactive-state) (**Figure12A-C**). These kinetic metastable states have transition from active to inactive through intermediate structure (transition-state) and vice-versa.^31^ The dynamic states would proceed through mechanistic pathways to initiate signaling process as expanding or compressing of the activation loop, outward/inward rotation of αC-helix and extended movements in the P-loop. It is clearly visible that the cavity of active site is enhanced in the presence of inhibitor to broken the R-spine obtained by moving apart of Glu residue on αC-helix and Phe residue in DFG motif associated in activation loop. The uncertainty of migrated residues could be withheld in particular state of kinase domain vertically from N-lobe towards C-lobe. In both states, the R-spine is broken differently in situ with all four residues moved apart. In the active state model kinase, the hydrophobic surface is broken between αC-helix bound Met-589 and DFG motif bound Phe-691 residues due to the extend space of activation loop and inward rotation of αC-helix. Inactive state model kinase consists β4-strand sheet (P-loop) bound Leu-620 and αC-helix bound Met-589 in situ R-spine opened in a discontinuous manner due to the outward rotation of αC-helix and the activation loop has recoiled into α-helix where DFG motif and αC-helix moves away from the P-loop of β-sheet (**Figure 13A-B**).

**Figure 13:**
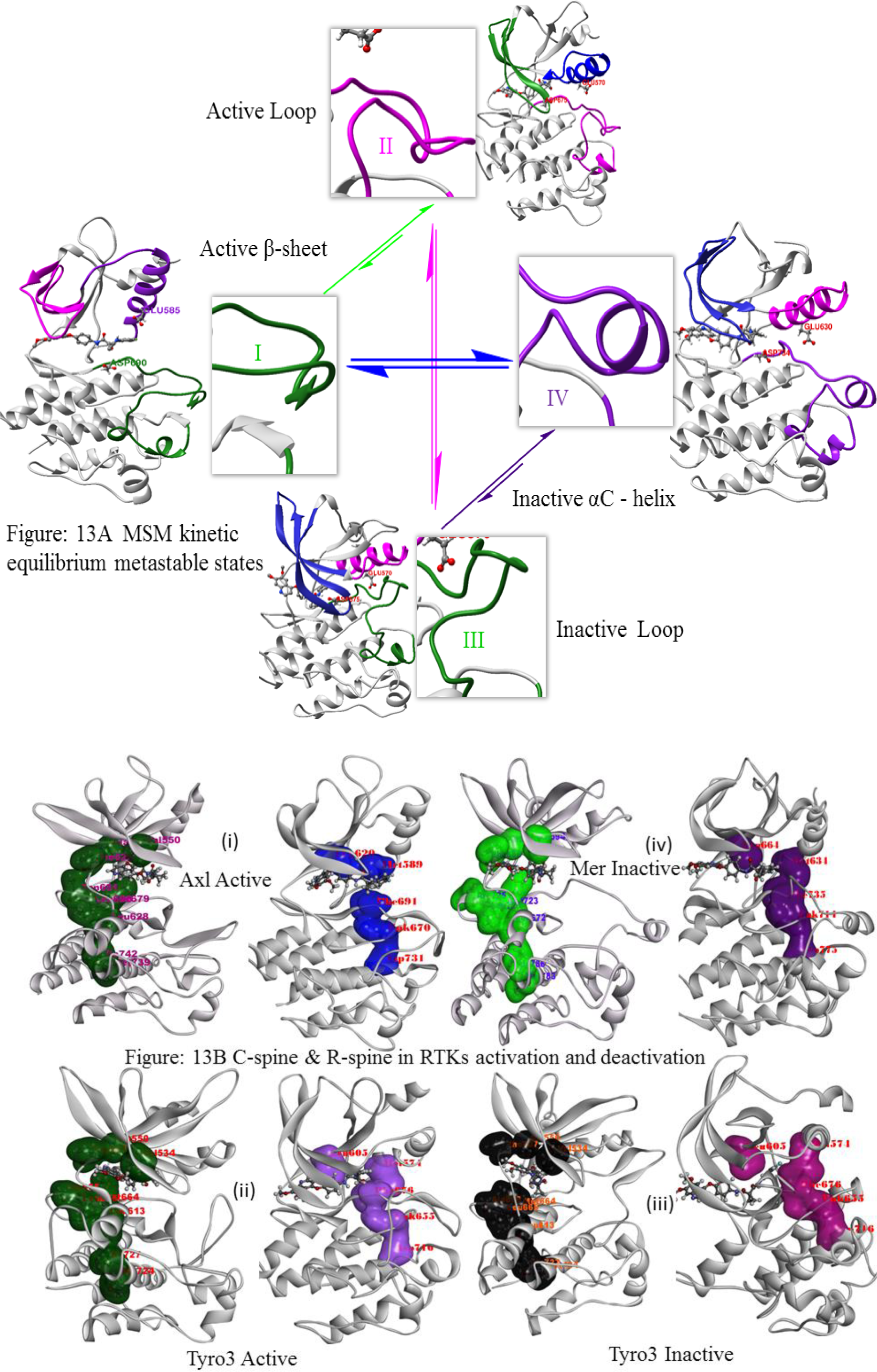
(13A) Schematic view of three kinetic equilibrium metastable states among HMM states involved allosteric activation and deactivation done in RTKs.(I) active state (II & III) transition states (IV) inactive state (13B) three kinetic equilibrium transition state have Catalytic (C) and Regulatory (R) spine mechanism while bound Cabozantinib in different states of RTKs from 1μs MD Simulations.

The hydrophobic surface is in a closed manner and continuous in apo form of active Tyro3, Axl and Mer RTKs. This space has expanded in the case of active state rather than inactive state model. This is achieved based upon activation loop refolded into β-sheet and π-stacking with β-sheet structure of catalytic loop in active state model (observed in Axl). The overall kinase bound drug should be propagated to its stationary distribution equal in Tyro3 active and inactive (Equal distribution). However, Axl and Mer kinase has higher stationary distribution with their active states (1.0 and 0.5 more as respectively) rather than inactive MSM states. It is concluded that the presence of inhibitor bound kinase domain proceeds as inactive state to block transduction of cellular mechanistic signal pathways in cancer therapy. RTKs are a major class of kinases involved in regular cellular metabolic activities for human physiological processes. Abnormal activation and overexpression of RTKs results in several forms of cancers. From one µs MD simulations each of apo, cabozantinib inhibitor bound active and inactive TAM kinases, we revealed metastable conformations between active and inactive sates. The location of R-spine and C-spine are mapped on the structures of TAM kinases. The cabozantinib binding stabilised the hidden markov state structures of active and inactive Axl, whereas the hidden markov state conformations from the three Mer structures are closely associated with each other. The αC-helical region is highly distinguished and its conformational flexibility is complementary to that of the activation loop. The R-spine is intact and vertically aligned in all apo TAM kinases. However, it is broken in cabozantinib bound active and inactive TAM kinases due to the expansion of protein core due to fluctuations in P-loop, αC- helix and activation loop. These changes also result in the loss of conserved salt bridge between the acidic residue on αC-helix and the basic residue in the activation loop. The dynamic movement of the R and C - spine residues are a result of the coordinated alterations in the kinase structural domain during the cellular signal transduction process. The selective kinase inhibitor (cabozantinib) arrests the activity of these overexpressed kinase domains via the dynamic movement of both these spines and distancing the space between αC-helix and activation-loop (**Figure S1b**). This can be supported from the results of distance plots shown in the active states of Tyro3, Axl and Mer that have undergone large expansion of protein core between αC-helix- A-loop in regulatory active site (**Figure 6a Column 2**). Therefore, the kinase activation done by most of active state mode.

From PCA, it is revealed that all TAM RTKs have random distribution of states. The analysis of metastable kinetic state forms revealed higher numbers of active state models in Axl and Mer RTKs than the number of kinetic transition states of inactive forms, however Tyro3 RTK has similar numbers of co-existed metastable state transitions among the active and inactive forms. The MSMs of the TAM kinases is expressed as different relaxation time scale intervals. The Tyro3 apo, Axl inactive and Mer active have higher relaxation time scales. The Tyro3 has equal contribution in active and inactive stationary distributions among kinetic HMM states. Five state kinetic metastable models were designed based upon active space distribution of HMMs of TAM RTKs. The apo Axl, inactive Mer and inactive Tyro3 have higher transition energies. These kinetic states are further validated with five MSM system to emphasize the hidden markov dynamic state analysis. Among the nine kinetic metastable states there are three HMM states classified as non-equilibrium kinetic transition states (Tyro3-apo S_3_-S_5_, Axl-inactive S_2_-S_5_ and Mer-active S_2_-S_4_) due to different kinetic transitions occurring among them (**Table-1**). The rest of six HMM states undergo S_4_-S_5_ kinetic transitions among five state model system. The activation loop undergoes β-sheet formation in the case of active Axl and α-helix formation in the case Mer inactive during S_4_-S_5_ kinetic transitions. In the case of Tyro3 active and inactive states, the activation loop remains in a random loop conformation. In summary, we observed salient changes in the spatial conformational states due to inhibitor binding to the active site during MD simulations in various regions of Tyro3, Axl and Mer kinases. From these research findings, the kinetic active and inactive state mechanism could explain how the inhibitor arrests the overexpressed RTKs in malignant cells, a key step to inhibit the kinase signaling pathway in cellular signaling process.

## Computational methods

### Structures of apo, active and inactive TAM RTK kinase domains

The three-dimensional crystal structures of Axl (PDB id: 5U6B) ^6d^ A and B chains were considered for inactive and active structures, respectively. The missing residues in the activation loop were built using “Model/Refine Loops” in “Structure Editing” tool in UCSF Chimera 1.12.^32^. The active and inactive homology model structures of Mer and Tyro3 were built based on the crystal structures of 5U6B B and A chains, respectively, using MODELLER as described previously. ^33, 34, 35a^

### Molecular docking of cabozantinib

The coordinates of the inhibitor, Foretinib were taken from the crystal structure (PDB id: 5IA4) and was used to deduce the structure cabozantinib.^36^ AutoDock, a molecular docking tool ^37^ was used to dock cabozantinib into the ATP binding pocket of the active and inactive conformers of TAM kinases as described previously.^35b^ The conformation with lowest binding energy and maximum docking poses was utilized for further MD simulations to decipher the molecular basis for interactions between protein and inhibitor.

### MD simulations

All MD simulations were achieved using AMBER ^38^ version 18.14 for the apo, active and inactive states of Tyro3, Axl and Mer kinase domain- cabozantinib complexes. The best docking pose of each complex was utilized as input for MD simulations. The force field charges for the entire systems were generated with Antechamber using am1bcc method.^39a, 40^ All input parameter files for MD simulations were generated after adding hydrogen atoms in tLEaP module using AMBER tools (ver.18.14) suite.^41^ Sodium and chloride ions were added to the systems to neutralize the charge, each molecular system was solvated within a 10 Å size box. The final ionic concentration for the systems was set to 100 mM. The Amberff99sb-idln force field was used for entire model system with TIP3P water model for AMBER molecular parameters.^42, 43^

All MD simulations were run at system temperature 300 K and 1 atm pressure with Monte Carlo barostat.^44^ Energy minimization was carried out by using steepest descent method around 40,000 cycles to overcome short range null contacts among the molecular system in solvent.^39b^ Long range electrostatic interactions were considered with Particle Mesh Ewald algorithm^45^ with cut-off range 9 Å and order 4. All model systems were equilibrated for 5 ns before the production run, and the coordinates in the production run were saved after every 5 ps.^46, 47^ The MD simulations of each molecular system was carried out for 1 µs MD simulations, accounting to a total of 9 µs simulations time.

### Post-MD data analysis

The MD trajectory data analysis was carried out using cpptraj and pytraj in Amber tools 18.^48^ The average structures after MD simulations, root mean square deviation (RMSD), root mean square fluctuation (RMSF) and principal component analysis (PCA) were derived from the trajectory analysis with various coding platforms provided by AMBER18.^49^ For the sake of data space minimization during post MD data analysis, the Markov State Model (MSM) analysis was carried out on 40K frames out of 200K frames and the PCA was carried out on data from 1K frames generated from each molecular system.

### Markov state model

To build the MSM, datasets of close accessible kinetic metastable states associated with protein conformational ensemble obtained from large scale simulations are required. These states can be defined in pyEMMA Python library as definite torsions, positions and distances of protein Cα data ^50^ to generate the MSM state models 40K conformations were sampled. All nine MD simulations datasets were transformed in terms of protein Cα backbone and dihedrals, distances of torsions, from raw MD datasets.^51^ All MD simulations trajectories were analysed for 200 ns (800 to 1000 ns) data by sampling the MSM predictions.^52a^ In the course of fast analysis, all nine MD simulations datasets (for 200 ns each) are carried out for subsample analysis to design and validate metastable models. These trajectory analyses identified kinetically metastable transitions among cluster k-means lag time of protein conformations.^53^ Python loaded libraries such as Interactive Python, standard Bio-python and Scikit-learn, PyEMMA^54–56^ were utilized for transformation of datasets in MSM model predictions. The extrapolation of the real time data into pictorial and graphic vectorised data points was achieved with matplotlib, and numpy data frames into 2D plotting space. The state distributions of kinetic metastable data points were featurized and cluster analysis was applied using time lagged independent component analysis (tICA). ^57^

### Data availability

All 9 µs MD structures and trajectory data, markov models are available upon reasonable request to corresponding author. All rest of the data held with manuscript supporting information including videos.

## Supporting information

This article contains supporting information.^58^

## Author information

### Corresponding Author

Dr. Lalitha Guruprasad, Ph.D. Professor School of Chemistry University of Hyderabad Hyderabad 500046 India. https://orcid.org/0000-0003-1878-6446

### Author

GATTA KRS NARESH Research Scholar School of Chemistry University of Hyderabad Hyderabad 500046 India. https://orcid.org/0000-0001-5194-0148

## Acknowledgements

NGKRS thanks University of Hyderabad for UGC Non-NET fellowship. The authors thanks DST-PURSE and UGC UPE2 for funding and CMSD for computational facilities.

## Conflicts of interest

The authors declare no competing interests.

## Abbreviations

ANM: Anisotropic normal modes
C-spine: Catalytic spine
DFG: Asp-Phe-Gly
EGF: Epidermal growth factor
HMM: Hidden Markov state models
JAK: Janus kinase
MEK/MAP2K: Mitogen-activated protein kinase
MSM: Markov’s state model
NumPy: Numerical Python
PCA: Principal component analysis
PCCA: Perron-cluster cluster analysis
PI3K: Phoshatidylinositol 3-kinase
PyEMMA: Python Emma’s Markov Model Algorithms
R-spine: Regulatory spine
RIN: Residue interaction network
RMSD: Root mean square deviation
RMSF: Root mean square fluctuation
RTKs: Receptor tyrosine kinases
SciPy: Scientific Python
TAM: Tyro3, Mer, Axl

